# α-integrins dictate distinct modes of type IV collagen recruitment to basement membranes

**DOI:** 10.1101/568964

**Authors:** Ranjay Jayadev, Qiuyi Chi, Daniel P. Keeley, Eric L. Hastie, David R. Sherwood

## Abstract

Basement membranes (BMs) are cell-associated extracellular matrices that support tissue integrity, signaling, and barrier properties. Type IV collagen is critical for BM mechanical and signaling functions, yet how it is directed into BMs *in vivo* is unclear. Through live-cell imaging of endogenous localization, conditional knockdown, and misexpression experiments, we uncovered distinct mechanisms of integrin-mediated collagen recruitment to *Caenorhabditis elegans* gonadal and pharyngeal BMs. The putative laminin-binding αINA-1/βPAT-3 integrin was selectively activated in the gonad and recruited laminin, which directed moderate collagen incorporation. In contrast, the putative Arg-Gly-Asp (RGD)-binding αPAT-2/βPAT-3 integrin was activated in the pharynx and recruited high levels of collagen independent of laminin. Through an RNAi screen, we further identified the small GTPase RAP-3 (Rap1) as a pharyngeal-specific PAT-2/PAT-3 activator that modulates collagen levels. Together, these studies demonstrate that tissues can use distinct mechanisms to direct collagen incorporation into BMs to precisely control collagen levels and construct diverse BMs.

## Introduction

Basement membrane (BM) is a thin, dense, sheet of extracellular matrix that underlies epithelia, endothelia, and surrounds most other tissues^1^. BMs are comprised primarily of two independent self-assembling scaffolds of laminin and type IV collagen. The laminin and type IV collagen networks are thought to associate through several cross-bridging molecules, including nidogen, perlecan, and agrin^2^. BMs are highly diverse, and this diversity arises from alternative splicing, post-translational modifications, and varying amounts of core BM components as well as numerous regulatory proteins that associate with BMs such as matricellular proteins, proteases, and growth factors^3,4^. The diversity in BMs regulates key cell and tissue properties, including cell polarity, cell differentiation, cell survival, tissue shaping, filtration, and resistance to mechanical stresses^5^. How diverse BMs are constructed on tissues is not well understood, particularly as many BM components are expressed and secreted from distant sources and selectively acquired from the extracellular fluid^3,6^.

Type IV collagen is highly conserved among animals, and its origination in unicellular ancestors was thought to have been a requirement for the formation of BMs and appearance of complex tissues in animals^7,8^. Type IV collagen is a heterotrimer made up of two α1 and one α2 chains that wind around each other into a long and rigid 400nm triple helix. The stiff triple helical structure together with covalent cross-linking between type IV collagen molecules bestows BMs mechanical resistance to tensile forces^7^. In addition to providing structural integrity, distinct tissue levels, unique modes of deposition, and targeted remodeling of collagen helps shape organs^9–13^. Highlighting its importance to human health, mutations in type IV collagen are associated with at least ten distinct genetic disorders that disrupt brain, kidney, muscle, and vascular tissues^7^. In mice, *Drosophila,* and *C. elegans*, type IV collagen is often secreted by cells into the extracellular space and then incorporated into epithelial BMs at distant sites^14–16^. Work in *Drosophila* and mouse embryos have indicated type IV collagen recruitment to BMs requires the presence of laminin^9,17,18^. Cell culture work has suggested that laminin deposition onto cell surfaces is mediated through interactions with integrin and dystroglycan receptors, and with sulfated glycolipids^2,19,20^. The cell surface mechanisms that mediate laminin and collagen deposition *in vivo*, however, remain elusive. Further, while most studies have examined initial BM formation in the embryo, little is known about BM construction on growing tissues, where most BMs are found.

Many factors have limited the study of how BMs are formed and the mechanisms by which type IV collagen is deposited into them. Laminin and type IV collagen are essential for embryonic animal development, limiting post-embryonic studies. Further, in vertebrates there are three distinct type IV collagens, 16 laminins, and at least 24 integrin αβ heterodimer receptors, making interpretation of genetic analysis daunting^6^. In addition, BM components and receptors have not yet been tagged in vertebrates with genetically encoded fluorescent proteins to dynamically examine their localization^21^. *C. elegans* is a powerful model system to address many of these experimental limitations. Type IV collagen and laminin, as well as many BM-interacting receptors, have been functionally tagged with fluorescent proteins, thus allowing dynamic imaging of BM *in vivo*^22–24^. Further, core BM matrix components and receptors are conserved in the worm but have not undergone the dramatic gene family expansion found in vertebrates. For example, *C. elegans* has only a single type IV collagen, two laminins, and two integrin heterodimers. Finally, RNAi can be used to conditionally knock down BM components post-embryonically to avoid early lethality, thus expanding the diversity of BMs that can be experimentally studied^6^.

In this work we have examined the mechanism of type IV collagen addition into the growing BMs covering two organs during *C. elegans* larval development—the pharynx, a rigid pumping feeding organ; and the gonad, a flexible cylindrical organ that houses the germline. Using RNAi-mediated knockdown and an mCherry-tagged type IV collagen reporter, we found that collagen addition to BM was required to maintain both organs’ structure, and that the pharyngeal BM contained significantly higher levels of collagen. Strikingly, while laminin was necessary for type IV collagen addition into the gonadal BM, collagen addition was independent of laminin in the pharyngeal BM. Through RNAi screening and examination of endogenously tagged BM-binding receptors, we discovered that the putative laminin-binding integrin heterodimer αINA-1/βPAT-3 mediated laminin and collagen addition to the gonadal BM, while the putative Arg-Gly-Asp tripeptide (RGD)^25^-binding integrin αPAT-2/βPAT-3 promoted collagen addition to the pharyngeal BM independent of laminin. However, both integrin heterodimers were expressed in the gonad and the pharynx, suggesting that their selective activation was controlled by tissue-specific effectors. Consistent with this hypothesis, domain-swapping experiments demonstrated that the intracellular domain of the α integrin PAT-2 drove BM recruiting activity of the INA-1 extracellular domain in the pharynx, where INA-1 is normally not active. Further, using an RNAi screen we identified RAP-3, an ortholog of the mammalian integrin-activating small GTPase Rap1, as a pharyngeal-specific activator of PAT-2/PAT-3. Together, these results identify integrin receptors as key *in vivo* mediators of collagen incorporation into BM during tissue growth and show that BM diversity and precise modulation of BM collagen levels can be driven in part by the tissue-specific activation of distinct integrins.

## Results

### The pharynx and gonad are BM-encased growing organs supported by type IV collagen

In *C. elegans*, collagen IV is a heterotrimer formed from two α1-like and one α2-like chains, encoded by the *emb-9* and *let-2* genes respectively^26^. Collagen is predominantly synthesized in body wall muscles, secreted into the extracellular space, and then recruited to the BM of other tissues^14,27^. To understand how type IV collagen is targeted to the BM of growing organs, we examined the BMs of the pharynx and the gonad (Figure 1A). The *C. elegans* pharynx is a rigid, contractile feeding apparatus largely composed of radially arranged muscle and marginal cells that form an epithelium^28^. The gonad is a flexible cylindrical reproductive organ, which is enwrapped predominantly by thin gonadal sheath cells^29^. The pharyngeal epithelium and gonadal sheath cells are both surrounded by BMs that support each organ^30^. Using a functional type IV collagen reporter (EMB-9∷mCherry^23^, referred to as collagen∷mCh) we examined surface area projections of the gonadal and pharyngeal BMs and found that the pharyngeal BM grew approximately three-fold in surface area during larval development (L1 through young adult), while the gonadal surface area increased over 90-fold (Figure 1B). To determine if type IV collagen addition was required to maintain BM and tissue integrity during pharynx and gonad growth, we depleted EMB-9 (α1-like chain) by RNAi beginning at the L1 larval stage and analyzed both organs at adulthood (72h, Figure 1A). We found that reduction of collagen frequently resulted in deformation of the anterior pharyngeal bulb (n=15/20 adult animals, Figure 1C), and severe distortion and rupturing of gonadal tissue (n=20/20 adult animals, Figure 1C). Similar results were observed upon knockdown of *let-2* (the α2 chain of collagen IV, n=20/20 animals examined). Linescan analysis of mean fluorescence intensity revealed a ~70% reduction in collagen∷mCh levels in the pharyngeal BM. Further, gaps in collagen signal appeared at regions of pharyngeal bulb deformation (Figure 1C). Collagen was reduced by ~60% in the gonad at the early L3 stage, prior to BM rupturing and the gonadal tissue becoming disorganized (Figure 1C). Taken together, these results indicate that addition of type IV collagen to the BM is necessary to maintain tissue integrity during growth of the pharynx and gonad.

**Figure 1.**
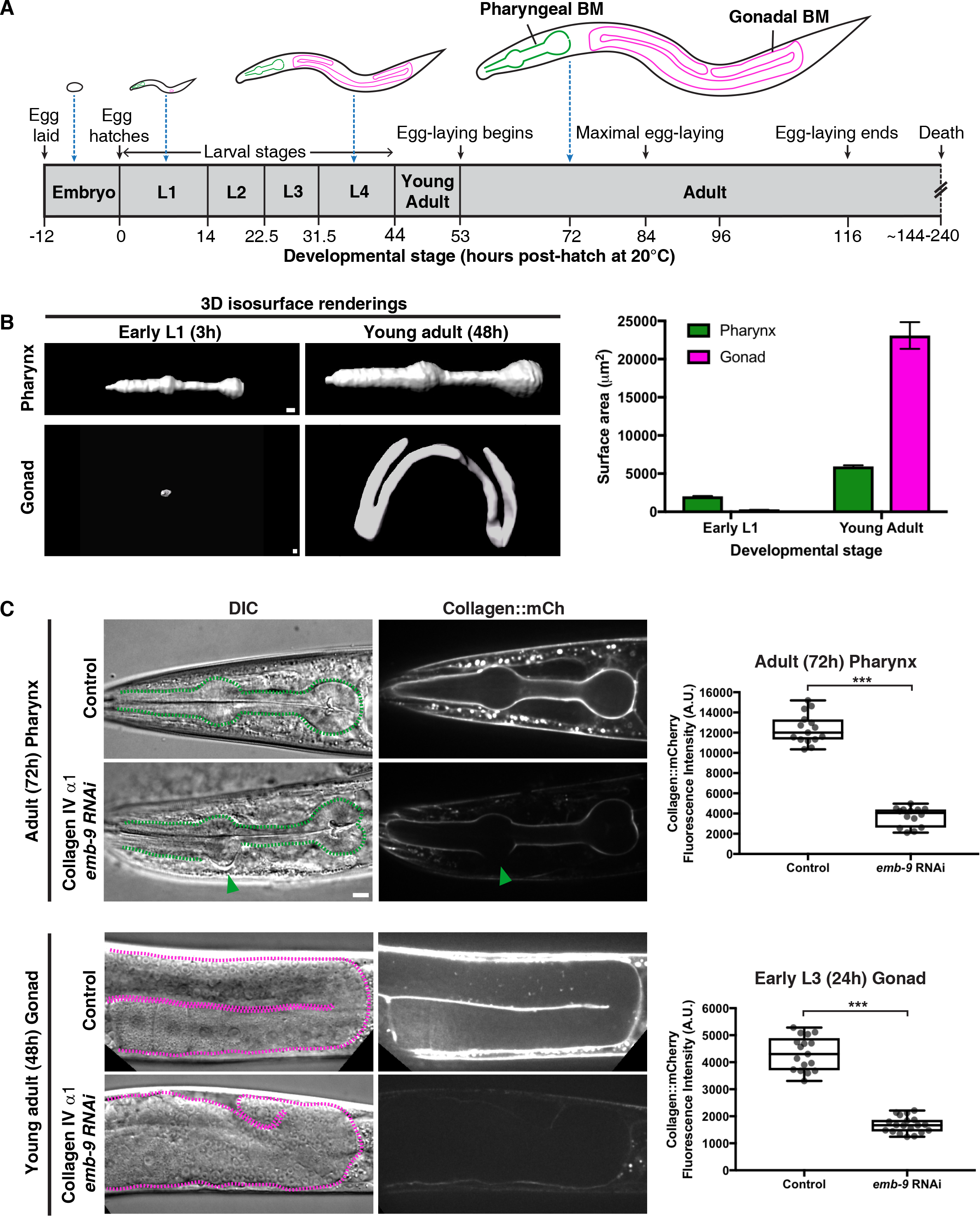
Basement membrane (BM) type IV collagen supports the growing *C. elegans* pharynx and gonad. **(A)** Illustrations of *C. elegans* at various stages of development (scaled to the length of the egg), with the pharyngeal and gonadal BMs outlined in green and magenta, respectively. **(B)** 3D isosurface renderings of pharyngeal and gonadal type IV collagen∷mCh at the early L1 versus young adult stages on the left, and quantification of surface area on the right. Bar graphs show mean surface area and error bars represent standard error of the mean (n=10 all stages). **(C)** DIC and collagen∷mCh fluorescence images of adult pharynxes (top left) and gonads (bottom left) of control (L4440 empty vector) and collagen IV α1 *(emb-9)* RNAi-treated 72h adult animals. Pharynxes in DIC images (top left) are outlined with green dotted lines. The green arrowhead indicates a deformation in the anterior pharyngeal bulb, corresponding to a region with undetectable collagen∷mCh signal. Mean pharyngeal BM collagen∷mCh fluorescence intensity in control (n=15) and *emb-9* RNAi-treated (n=14) 72h adult animals are quantified on the top right. Dotted magenta lines outline gonads in DIC images (bottom left). The gonads of *emb-9 RNAi*-treated animals are severely misshapen, correlating with near undetectable BM collagen∷mCh signal. Mean gonadal BM collagen∷mCh fluorescence intensity in control (n=17) and *emb-9* RNAi-treated (n=19) early L3 animals are quantified on the bottom right. \*\*\**p*<0.0001, unpaired two-tailed Student’s *t* test. Box edges in boxplots depict the 25^th^ and 75^th^ percentile, the line in the box indicates the median value, and whiskers mark the minimum and maximum values. Scale bars are 10μm.

### The pharyngeal BM is collagen-rich, but laminin-poor, compared to the gonadal BM

BMs of different tissues often vary in levels and composition of matrix components^31,32^. To determine if type IV collagen levels differ between the gonadal and pharyngeal BMs, we quantified collagen∷mCh fluorescence intensity in both organs and found that collagen was present at two-fold higher levels in the pharyngeal BM versus the gonadal BM in 72h adults (Figure 2). As embryonic studies have suggested that collagen is recruited to BMs through association with laminin networks^9,17,18,30^, we next asked if higher levels of laminin might be required to recruit more collagen to the pharyngeal BM. Laminins are heterotrimers comprised of an α, β, and γ chain. *C. elegans* encodes two α subunits (*lam-3* and *epi-1*), one β subunit (*lam-1*), and one γ subunit (*lam-2*), that form two distinct laminin heterotrimers^26^. To assess laminin levels in the BM, we used CRISPR/Cas9 to encode *mNeonGreen* at the *lam-2* genomic locus, generating worms expressing the LAM-2∷mNeonGreen fusion protein (referred to as laminin∷mNG). Surprisingly, we found that there were two-fold lower levels of laminin in the pharyngeal BM compared to the gonadal BM (Figure 2). Thus, there are higher levels of type IV collagen in the pharyngeal BM compared to the gonadal BM, but this collagen enrichment is not correlated with increased laminin levels.

**Figure 2.**
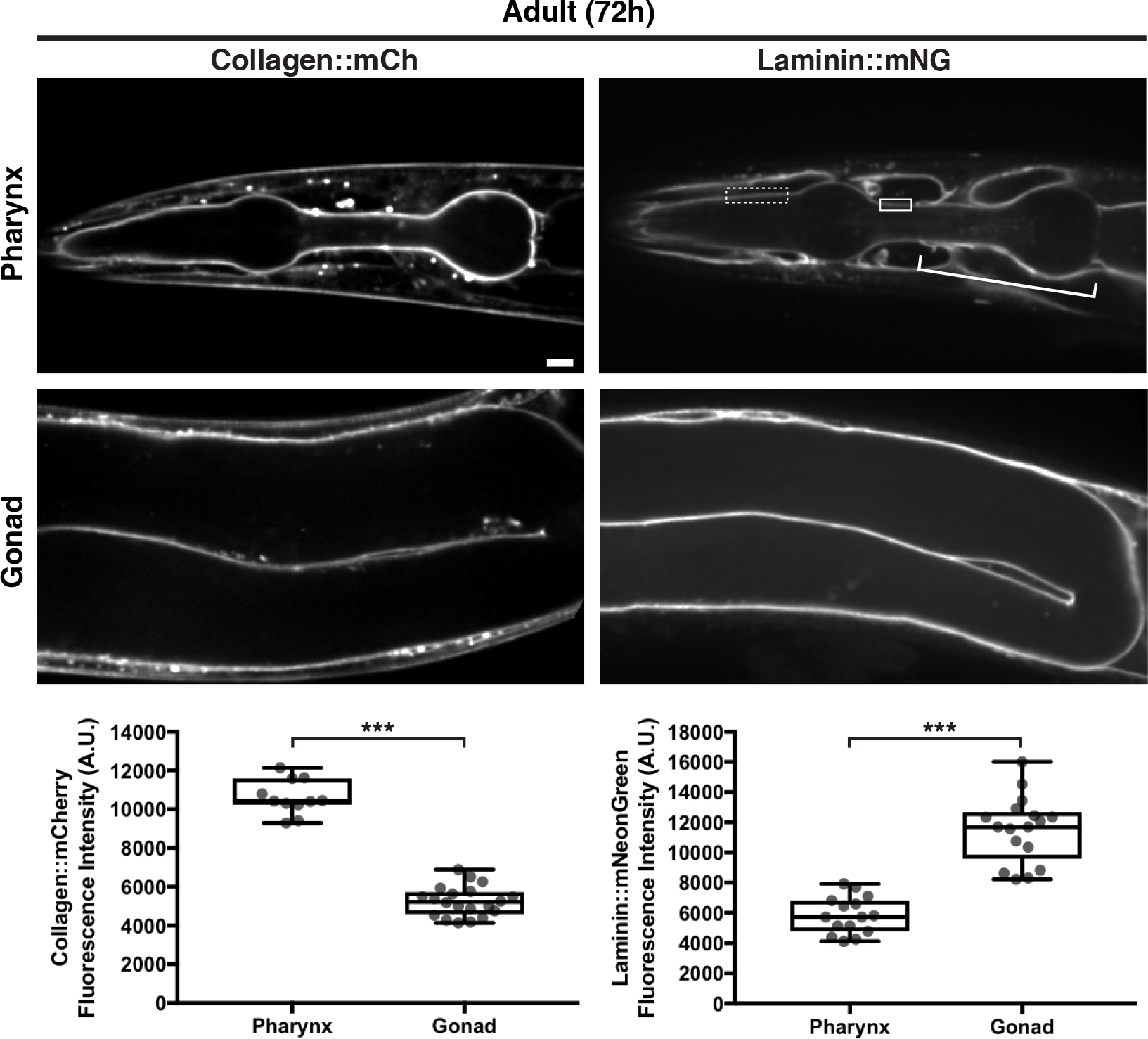
The pharyngeal BM is collagen rich, while the gonadal BM is laminin-rich. Fluorescence images of collagen∷mCh and laminin∷mNG in the pharynx and gonad of 72h adult animals. As laminin∷mNG localizes strongly to the nerve ring BM that surrounds and contacts the pharyngeal BM at several regions (bracket), we confined our measurements of laminin signal to the dotted box where clear pharyngeal BM laminin is visible in this and all relevant figures, except figure 6C, where laminin signal in the solid box was also measured. Quantification of collagen∷mCh and laminin∷mNG levels in the pharyngeal (collagen∷mCh n=11; laminin∷mNG n=15) and gonadal BMs (collagen∷mCh n=20; laminin∷mNG n=17) are shown in boxplots at the bottom. *** *p*<0.0001, unpaired two-tailed Student’s *t* test. Box edges in boxplots depict the 25^th^ and 75^th^ percentile, the line in the box indicates the median value, and whiskers mark the minimum and maximum values. Scale bars are 10μm.

### Laminin is required for collagen recruitment to the gonadal but not pharyngeal BM

Our finding that the collagen enrichment in the pharyngeal BM relative to the gonadal BM is not correlated with similarly increased laminin levels suggested that the interaction between type IV collagen and laminin may differ in the BMs of these tissues. We next asked whether laminin was required to recruit collagen to these BMs. We used RNAi to deplete LAM-2 (the sole laminin γ subunit) beginning in the L1 larval stage and analyzed BM collagen levels during subsequent larval development and adulthood. By the early L3 larval stage (24h RNAi treatment), there was a ~50% reduction in gonadal BM laminin levels (Figure S1A), correlating with a similar reduction in type IV collagen levels (Figure 3A). Breaks in the BM were first observed shortly after this time (~26h RNAi treatment, n=12/20 animals), followed by BM rupturing (Mid L4, Figure 3A, n=20/20 worms), and ultimately gonadal tissue disintegration (72h adult, Figure 3A, n=20/20 animals examined). In contrast, the pharyngeal BM collagen levels were unaffected despite a strong knockdown of laminin (~70% reduction in laminin levels by 96h adult, Figure S1B), and there were no defects in pharyngeal structure (n=14/14 animals). These observations suggest that type IV collagen recruitment to the growing BM is distinct in each tissue—laminin-dependent in the gonad but laminin-independent in the pharynx.

**Figure 3.**
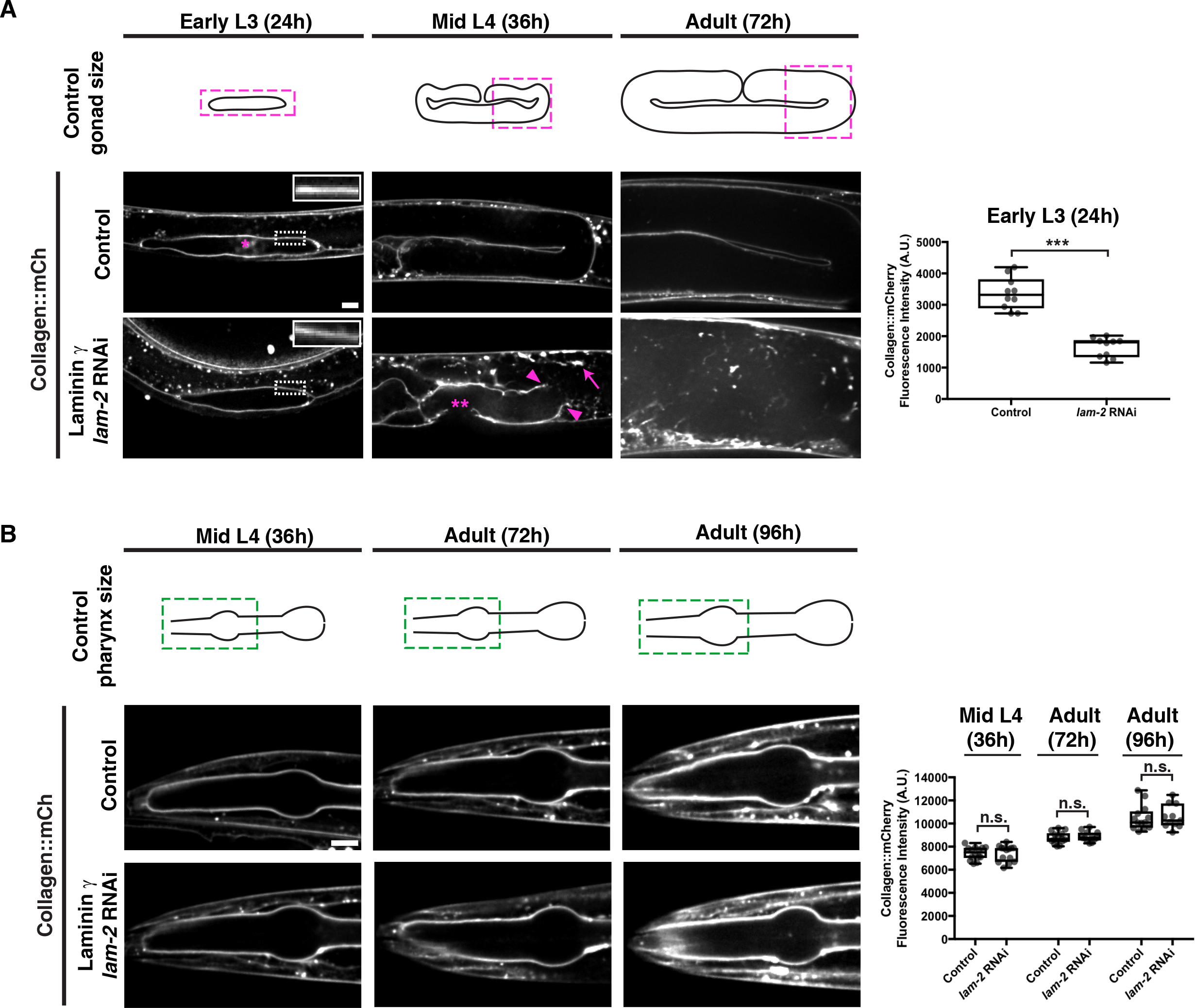
Collagen recruitment to the gonadal but not pharyngeal BM is dependent on laminin. **(A)** Fluorescence images of gonadal BM collagen∷mCh shown at the early L3, mid-L4, and 72h adult stages in control and laminin γ *(lam-2)* RNAi-treated animals (RNAi fed at the L1 stage onwards). Control gonad size at these stages are shown in schematics, and the magenta boxes denote regions of the gonad shown in the figure. Collagen∷mCh signal in the dotted box regions are magnified in insets. The asterisk indicates non-BM collagen∷mCh signal from coelomocytes. Gonadal BM collagen∷mCh fluorescence intensity in control (n=10) and *lam-2* (n=10) RNAi-treated early L3 animals is quantified on the right. By the mid-L4 stage, rupturing of the gonadal BM (magenta arrowheads) and abnormal collagen aggregation (magenta arrow) was observed in *lam-2* RNAi-treated animals (n=20/20). The double asterisks mark a break in the BM due to anchor cell invasion, a normal morphogenetic event during *C. elegans* vulval development. **(B)** Fluorescence images of pharyngeal BM collagen∷mCh in control and *lam-2* RNAi-treated animals shown at the mid L4, 72h adult, and 96h adult stages. Control pharynx size at these stages are shown in schematics, and the green boxes denote regions of the pharynx shown in the figure. Pharyngeal BM collagen∷mCh fluorescence intensity in control and *lam-2* RNAi-treated animals at all three stages are quantified on the right (mid L4 control n=14, *lam-2* RNAi n=14; 72h adult control n=14, *lam-2* RNAi n=12; 96h adult control n=13, *lam-2* RNAi n=11). *** *p*<0.0001, unpaired two-tailed Student’s *t* test; n.s., *p*>0.05. Box edges in boxplots depict the 25^th^ and 75^th^ percentile, the line in the box indicates the median value, and whiskers mark the minimum and maximum values. Scale bars are 10μm.

### INA-1/PAT-3 integrin mediates collagen recruitment to the gonadal BM in a laminin-dependent manner

Work in cell culture has suggested that interactions with laminin-binding integrin and dystroglycan receptors as well as cell surface sulfated glycolipids might seed laminin polymerization to initiate BM assembly, leading to subsequent collagen recruitment^19,33^. Because of possible redundancy between these mechanisms and within the large integrin gene family in vertebrates, however, there has not yet been clear genetic evidence for any of these mechanisms in regulating laminin and collagen recruitment to BMs *in vivo*^34^. As C*. elegans* does not synthesize sulfated glycolipids^35^, we hypothesized that tissue-specific roles for integrin and dystroglycan might explain the different modes of collagen recruitment to the pharyngeal and gonadal BMs.

To investigate the cell surface interactions that recruit type IV collagen to developing tissues, we initially focused on the gonadal BM. We first examined the dystroglycan receptor. *C. elegans* harbors a single vertebrate dystroglycan ortholog, *dgn-1*^36^. Endogenously tagged DGN-1∷mNG^37^ was expressed in the gonad and localized to the gonad-BM interface (Figure S2A). Knockdown of *dgn-1* by RNAi eliminated detectable DGN-1∷mNG but did not affect either laminin or collagen levels in adult animals (Figure S2B). We next investigated the integrin receptor system. *C. elegans* assembles two integrin heterodimers, composed of the α subunits PAT-2 (most similar to RGD-binding integrins) or INA-1 (most similar to laminin-binding integrins) dimerized to the sole β subunit PAT-3^26^. Endogenously tagged INA-1∷mNG, PAT-2∷2XmNG, and PAT-3∷mNG were expressed in the gonad and localized to the cell-BM interface (Figure 4A). L1 RNAi mediated knockdown of *ina-1* and *pat-3* each resulted in a ~50% reduction in collagen∷mCh levels in adult animals (Figure 4B), suggesting that the INA-1/PAT-3 integrin heterodimer is required for type IV collagen recruitment to the gonadal BM. Importantly, RNAi against *ina-1* and *pat-3* did not appear to alter collagen secretion from the muscle cells (Figure S3A), suggesting a direct collagen-recruiting function for INA-1/PAT-3 in the gonad. As gonadal BM collagen localization is dependent on laminin, we analyzed laminin∷mNG levels upon *ina-1* and *pat-3* knockdown and found a similar ~50% reduction (Figure 4B) in each condition. Surprisingly, depletion of the other integrin α subunit PAT-2 by RNAi yielded an increase in BM collagen and laminin levels (Figure 4B). We speculated that this could be due to more INA-1/PAT-3 heterodimers forming upon reduction of PAT-2 levels. Consistent with this idea, RNAi against *pat-2* more than doubled INA-1∷mNG levels at cell surfaces contacting the gonadal BM (Figure S3B).

**Figure 4.**
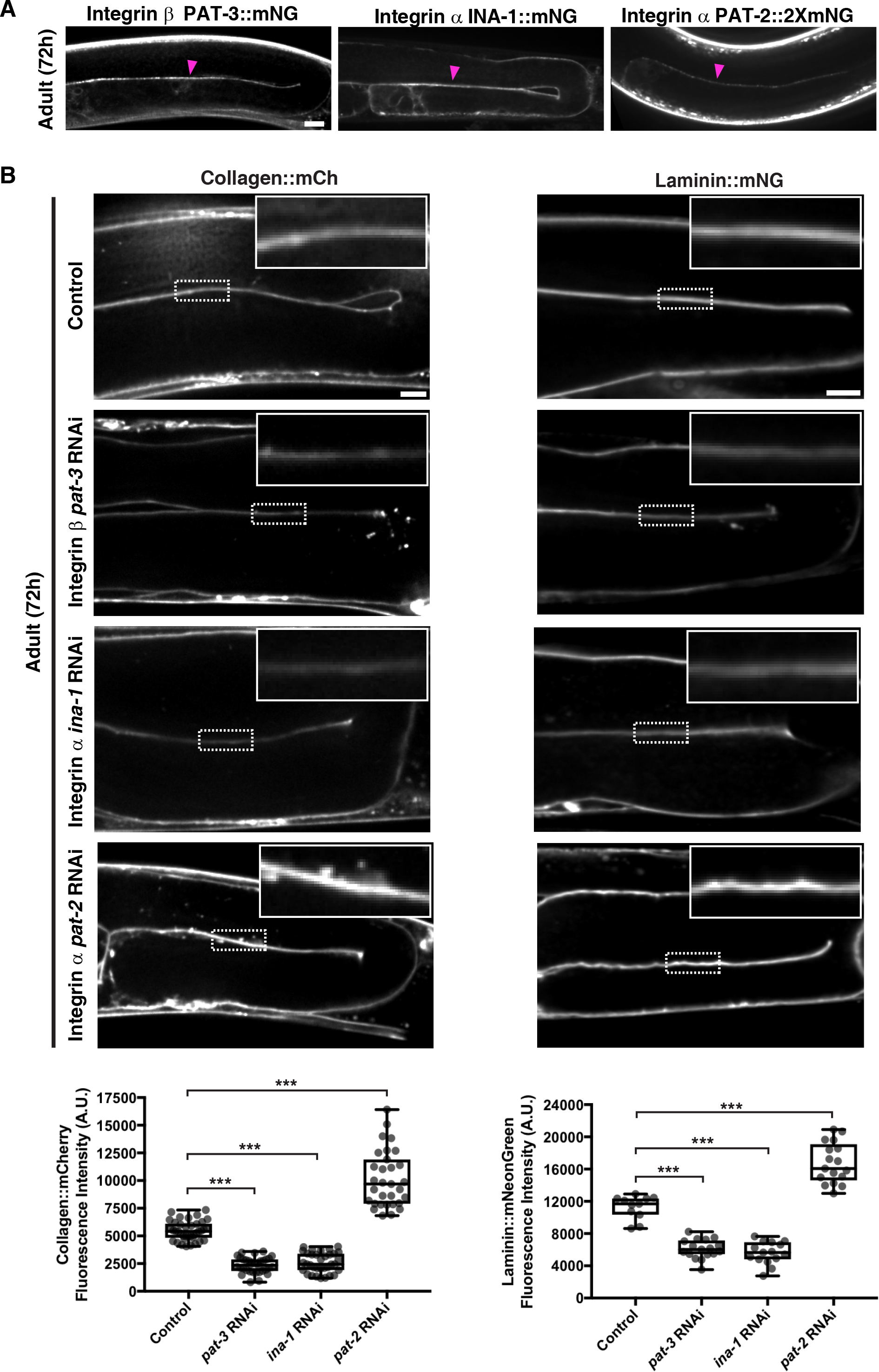
The INA-1/PAT-3 integrin heterodimer is required for laminin-dependent collagen recruitment to the gonadal BM. **(A)** Fluorescence images of the integrin β subunit PAT-3∷mNG and the integrin α subunits INA-1∷mNG and PAT-2∷2xmNG in the gonad. Magenta arrowheads indicate enrichment of fluorescence signal at the cell-BM interface. **(B)** Fluorescence images of gonadal BM collagen∷mCh (left) and laminin∷mNG (right) in control versus *pat-3*, *ina-1*, and *pat-2* RNAi-treated 72h adult animals. Regions of the BM in the dotted boxes are magnified in insets. Quantification of collagen∷mCh (control n=38; *pat-3* RNAi n=34; *ina-1* RNAi n=34; *pat-2* RNAi n=31) and laminin∷mNG (control n=11; *pat-3* RNAi n=17; *ina-1* RNAi n=15; *pat-2* RNAi n=17) fluorescence intensity for all treatments is shown at the bottom. *** *p*<0.0001, one-way analysis of variance (ANOVA) followed by post-hoc Dunnett’s test. Box edges in boxplots depict the 25^th^ and 75^th^ percentile, the line in the box indicates the median value, and whiskers mark the minimum and maximum values. Scale bars are 10μm.

To determine if any other matrix receptors contribute to collagen recruitment to the gonadal BM, we performed RNAi against worm orthologs of glypican (*gpn-1* and *lon-2*), LAR-RPTP (*ptp-3*), teneurin (*ten-1*), and discoidin domain receptors (*ddr-1* and *ddr-*2)^26^. Knockdown of these receptors did not affect gonadal collagen levels (Figure S3C). Taken together, our findings suggest that the putative laminin-binding INA-1/PAT-3 integrin heterodimer is the predominant cell surface receptor that mediates collagen IV recruitment to the gonadal BM through a laminin-dependent mechanism.

### The PAT-2/PAT-3 integrin heterodimer promotes collagen recruitment to the pharyngeal BM independent of laminin

We next sought to determine the cell surface receptor(s) mediating collagen recruitment to the pharyngeal BM. DGN-1∷mNG expression was not detectable in the pharynx (Figure S4A). In contrast, all three integrin subunits were present in the pharyngeal epithelium and localized to the pharynx-BM interface (Figure 5A). RNAi against *ina-1* did not affect pharyngeal BM collagen or laminin levels (96h RNAi treatment, Figure 5B). However, RNAi targeting *pat-2* and *pat-3* each reduced pharyngeal BM collagen levels by over 40% (Figure 5B). Notably, RNAi against *pat-2* or *pat-3* did not affect pharyngeal BM laminin levels (Figure 5B). These results suggest the PAT-2/PAT-3 integrin promotes pharyngeal BM collagen recruitment independent of laminin. To determine whether PAT-2 is sufficient to recruit collagen to the pharyngeal BM, we over-expressed PAT-2∷mNG specifically in pharyngeal muscle cells by using the *myo-2* promoter (Figure 5C). Mosaic analysis of adult animals revealed that pharyngeal BM collagen∷mCh levels were increased by ~30% in regions where PAT-2 was over-expressed (Figure 5C). Finally, RNAi-mediated reduction of other major matrix receptors did not affect collagen levels in the pharyngeal BM (Figure S4B). Together, these observations indicate that the putative RGD-binding PAT-2/PAT-3 integrin heterodimer is the predominant cell surface receptor that is required and sufficient to mediate type IV collagen recruitment to the pharyngeal BM independent of laminin. Further, these results suggest that the recruitment of laminin in the growing pharyngeal BM is not dependent on integrins (either INA-1/PAT-3 or PAT-2/PAT-3) but is mediated by an unknown mechanism.

**Figure 5.**
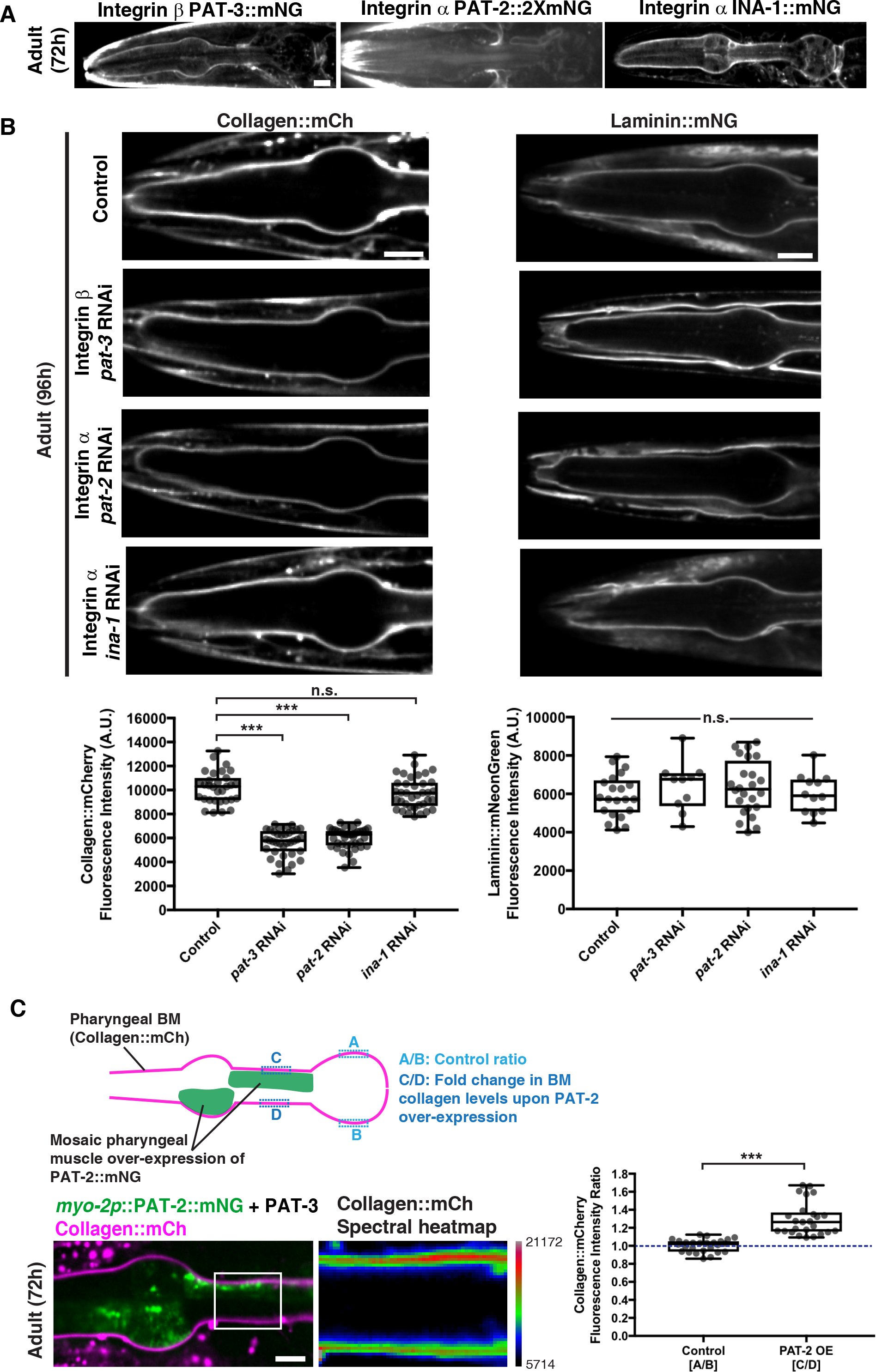
The PAT-2/PAT-3 integrin heterodimer is required for laminin-independent collagen recruitment to the pharyngeal BM. **(A)** Fluorescence images of the integrin β subunit PAT-3∷mNG and the integrin α subunits PAT-2∷2xmNG and INA-1∷mNG in the pharynx. **(B)** Fluorescence images of pharyngeal BM collagen∷mCh (left) and laminin∷mNG (right) in control versus *pat-3*, *pat-2*, and *ina-1* RNAi-treated 96h adult animals. Quantification of collagen∷mCh (control n=31; *pat-3* RNAi n=36; *pat-2* RNAi n=39; *ina-1* RNAi n=37) and laminin∷mNG (control n=21; *pat-3* RNAi n=10; *pat-2* RNAi n=24; *ina-1* RNAi n=12) fluorescence intensity for all treatments is shown at the bottom. *** *p*<0.0001, one-way ANOVA followed by post-hoc Dunnett’s test. n.s., *p*>0.05, one-way ANOVA. **(C)** A schematic outlining mosaic over-expression of PAT-2∷mNG in the pharyngeal muscle cells and method for quantification of collagen levels is shown on top. A merged fluorescence image of *myo-2p*∷PAT-2∷mNG (green) and collagen∷mCh (magenta) in a 72h adult pharynx is shown on the bottom left. A spectral heatmap of collagen∷mCh fluorescence intensity in the boxed region is shown on the bottom right. Quantification of the fold change in pharyngeal BM collagen∷mCh fluorescence intensity upon PAT-2 over-expression is shown on the right (n=25). *** *p*<0.0001, paired two-tailed Student’s *t*-test. Box edges in boxplots depict the 25^th^ and 75^th^ percentile, the line in the box indicates the median value, and whiskers mark the minimum and maximum values. Scale bars are 10μm.

### The PAT-2 intracellular domain dictates BM recruiting activity of integrins in the pharynx

All integrin subunits were expressed in both the gonadal and pharyngeal tissues. Thus, we next sought to determine the mechanism controlling the tissue-specific activity of the two integrin heterodimers in matrix recruitment. Recent studies have suggested that the diverse intracellular C-terminal membrane-distal (CTMD) regions of vertebrate α integrins may provide specificity for integrin inside-out activation—a form of integrin activation where intracellular regulators bind the cytoplasmic tails of integrin heterodimers and trigger conformational changes that allow high affinity binding of the integrin extracellular domain with matrix ligands^38^. As the sole difference between INA-1/PAT-3 and PAT-2/PAT-3 integrin heterodimers is the α subunit, we speculated that their tissue-specific activity in collagen recruitment could be regulated by their distinct α-integrin intracellular domains mediating inside-out activation. Indeed, INA-1 and PAT-2 diverge significantly in their CTMD regions (Figure 6A).

**Figure 6.**
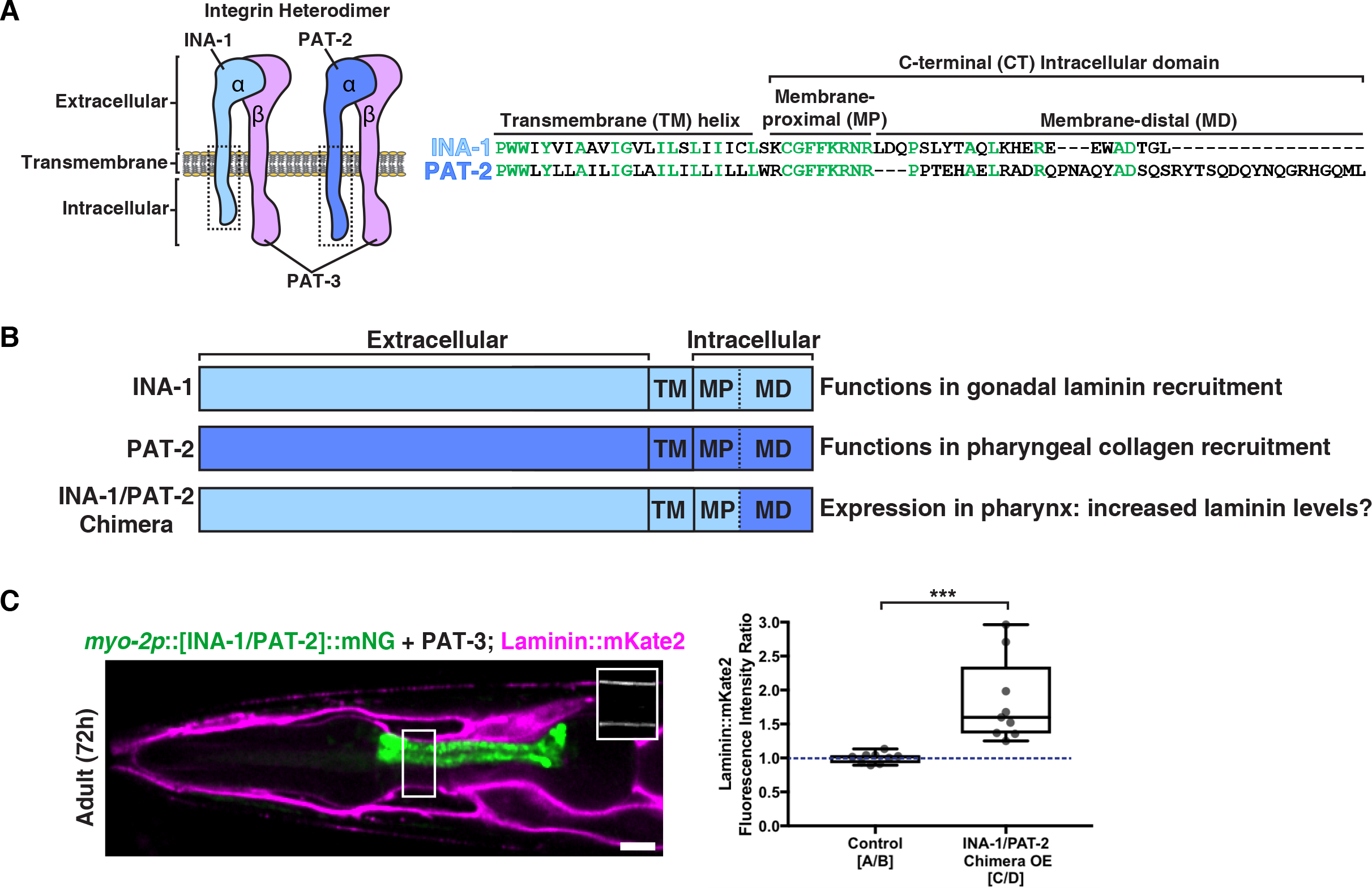
The intracellular domain of PAT-2 controls the activity of PAT-2/PAT-3 in the pharynx to promote collagen recruitment. **(A)** A schematic of the two integrin heterodimers expressed in the worm, highlighting the extracellular, transmembrane, and C-terminal intracellular regions of these proteins. Amino acid alignments of the boxed transmembrane and intracellular regions of INA-1 and PAT-2 are shown on the right. Highly conserved residues are shown in green. **(B)** Schematic of chimeric integrin α subunit. Determined functions of INA-1 and PAT-2 and the predicted outcome of INA-1/PAT-2 chimeric integrin expression in the pharynx are listed. **(C)** A merged fluorescence image of *myo-2p*∷[INA-1/PAT-2]∷mNG (green) and laminin∷mKate2 (magenta) in a 72h adult pharynx is shown. Laminin∷mKate2 signal in the boxed region is magnified in the inset. Quantification of the fold change in pharyngeal BM laminin∷mKate2 fluorescence intensity upon over-expression of the INA-1/PAT-2 chimera is shown on the right (n=9). *** *p*<0.0001, paired two-tailed Student’s *t*-test. Box edges in boxplots depict the 25^th^ and 75^th^ percentile, the line in the box indicates the median value, and whiskers mark the minimum and maximum values. Scale bars are 10μm.

Building off of these studies, and our findings that INA-1/PAT-3 integrin is expressed in the pharynx but does not recruit laminin to the pharyngeal BM, we hypothesized that the pharynx-active intracellular domain of PAT-2 fused to the laminin-binding extracellular domain of INA-1 might be able to recruit laminin in this tissue. We engineered animals that express this chimeric α integrin subunit in the pharynx (*myo-2p*∷INA-1[EX]∷PAT-2[CTMD]∷mNG, Figure 6B). Over-expression of this chimeric integrin resulted in a ~50% increase in laminin levels in regions of the BM contacting chimera-expressing pharyngeal cells (Figure 6C). Notably, as shown earlier (Figure 5B), full-length INA-1 does not normally recruit laminin to the larval pharyngeal BM. Together, our findings suggest that the cytoplasmic domain of PAT-2 facilitates the PAT-2/PAT-3-mediated recruitment of collagen to the pharyngeal BM, and when attached to the extracellular domain of INA-1, triggers laminin recruitment instead.

### RAP-3 is a pharyngeal-specific activator of PAT-2/PAT-3 integrin

Our results suggested that there might be tissue-specific factors that activate PAT-2/PAT-3 in the pharynx via the PAT-2 CTMD region to mediate BM matrix recruitment. Proteomic, localization, and genetic screening approaches have identified numerous proteins that localize with integrins or regulate integrin activity^39^. To identify potential tissue-specific regulators of PAT-2/PAT-3 integrin-mediated collagen IV recruitment to BM, we performed a targeted RNAi screen (Table S1) of *C. elegans* orthologs of 33 known integrin-associated or regulatory proteins^40–42^. We searched for proteins whose loss reduced type IV collagen levels in the pharyngeal BM but did not affect gonadal BM collagen levels. RNAi against one gene, *rap-3*, led to a ~40% reduction of pharyngeal type IV collagen levels without affecting gonadal BM collagen, suggesting it could be a specific activator of PAT-2/PAT-3 in the pharynx (Figures 7A and 7B).

**Figure 7.**
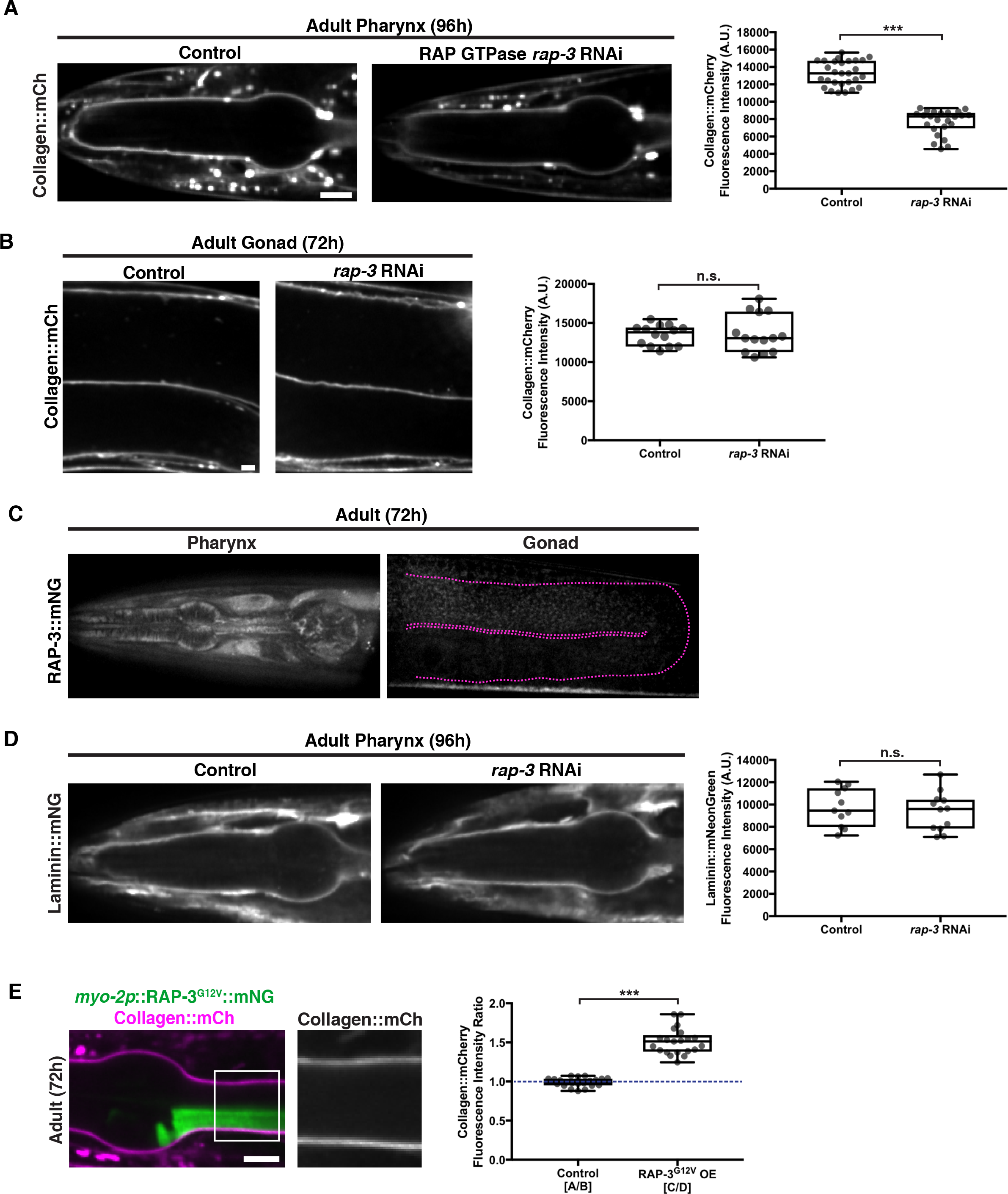
RAP-3 regulates the PAT-2/PAT-3-driven recruitment of collagen into the pharyngeal BM. **(A)** Fluorescence images of pharyngeal BM collagen∷mCh in control and *rap-3* RNAi-treated 96h adult animals, with quantification of collagen∷mCh fluorescence intensity on the right (control n=28; *rap-3* RNAi n=24). **(B)** Fluorescence images of gonadal BM collagen∷mCh in control and *rap-3* RNAi-treated 72h adult animals, with quantification of collagen∷mCh fluorescence intensity on the right (control n=14; *rap-3* RNAi n=14). **(C)** Fluorescence images of RAP-3∷mNG localization in the pharynx and its absence in the gonad (outlined in magenta). **(D)** Fluorescence images of pharyngeal BM laminin∷mNG in control and *rap-3* RNAi-treated 96h adult animals, with quantification of laminin∷mNG fluorescence intensity on the right (control n=11; *rap-3* RNAi n=12). *** *p*<0.0001, unpaired two-tailed Student’s *t* test; n.s., *p*>0.05. **(E)** Merged fluorescence image of *myo-2p*∷RAP-3^G12V^∷mNG (green) and collagen∷mCh (magenta) in a 72h adult pharynx. Pharyngeal BM collagen∷mCh signal in the boxed region is magnified on the right. Quantification of fold increase in collagen∷mCh fluorescence intensity upon RAP-3^G12V^ over-expression is shown on the right (n=21). *** *p*<0.0001, paired two-tailed Student’s *t*-test. Box edges in boxplots depict the 25^th^ and 75^th^ percentile, the line in the box indicates the median value, and whiskers mark the minimum and maximum values. Scale bars are 10μm.

The *rap-3* gene encodes a Rap-like protein most similar to the mammalian Rap1 isoforms Rap1A and Rap1B^43^. Rap1 is a member of the Ras family of small GTPases, and Rap1B regulates inside-out activation of integrins in blood cells, including platelets and lymphocytes, to facilitate their adhesion and migration^44–46^. To determine if the RAP-3 protein might be a tissue-specific regulator of PAT-2/PAT-3 integrin activity, we first generated RAP-3∷mNG-expressing animals using CRISPR/Cas9 genome editing. Consistent with a pharynx-specific function, we found that RAP-3∷mNG was expressed in the pharynx of larval and early adult animals, but was not detectable in gonadal tissue (Figure 7C and Figure S5A).

We hypothesized that if RAP-3 specifically activates PAT-2/PAT-3 to recruit type IV collagen, then it would not be involved in laminin recruitment. Supporting this notion, we found that pharyngeal BM laminin levels were unaffected by RNAi targeting *rap-3* (Figure 7D). Further, pharyngeal-specific over-expression of a constitutively active form of RAP-3 (RAP-3^G12V^)^47^ resulted in a ~50% increase in collagen levels in regions of the pharyngeal BM in contact with RAP-3^G12V^ over-expressing cells (Figure 7E). To test whether RAP-3 and PAT-2 might function in the same genetic pathway to mediate type IV collagen recruitment, we first generated a *rap-3* null mutation, *rap-3(qy67)*, using CRISPR/Cas9 genome editing. However, we were unable to view homozygous *rap-3(qy67)* mutant larval animals due to embryonic lethality. To circumvent embryonic lethality, we knocked both *pat-2* and *rap-3* down simultaneously by RNAi beginning at the L1 larval stage. The combined RNAi against *rap-3* and *pat-2* did not worsen the reduction in pharyngeal BM collagen levels caused by individual *pat-2* or *rap-3* knockdown, consistent with these genes functioning within the same pathway (Figure S5B). We also found that knockdown of *rap-3* did not alter PAT-2 or PAT-3 localization or levels (Figure S5C), suggesting that it does not activate PAT-2/PAT-3 by regulating integrin trafficking. Together, our observations support the idea that RAP-3 is a pharyngeal-specific activator of the PAT-2/PAT-3 integrin, triggering its ability to recruit type IV collagen to the pharyngeal BM.

Finally, we wanted to test if ectopic expression of RAP-3 in the gonad was sufficient to activate PAT-2/PAT-3 in this tissue to facilitate collagen recruitment. We used the *inx-8* promoter to drive RAP-3∷mNG expression in the gonadal sheath cells that contact BM^48^ (Figure 8A). Strikingly, we found that ectopic RAP-3 expression in the gonadal sheath cells increased collagen levels by ~80% in regions of the BM contacting RAP-3-expressing cells (Figure 8A). We predicted that if RAP-3 was triggering collagen recruitment by activating PAT-2/PAT-3 in the gonad, then laminin levels would not be altered. Consistent with this idea, we found that gonadal expression of RAP-3 did not affect BM laminin levels (Figure 8B). Together these observations strongly suggest that RAP-3 is a tissue-specific activator of PAT-2/PAT-3 integrin, which directs the laminin-independent addition of type IV collagen into the BM.

**Figure 8.**
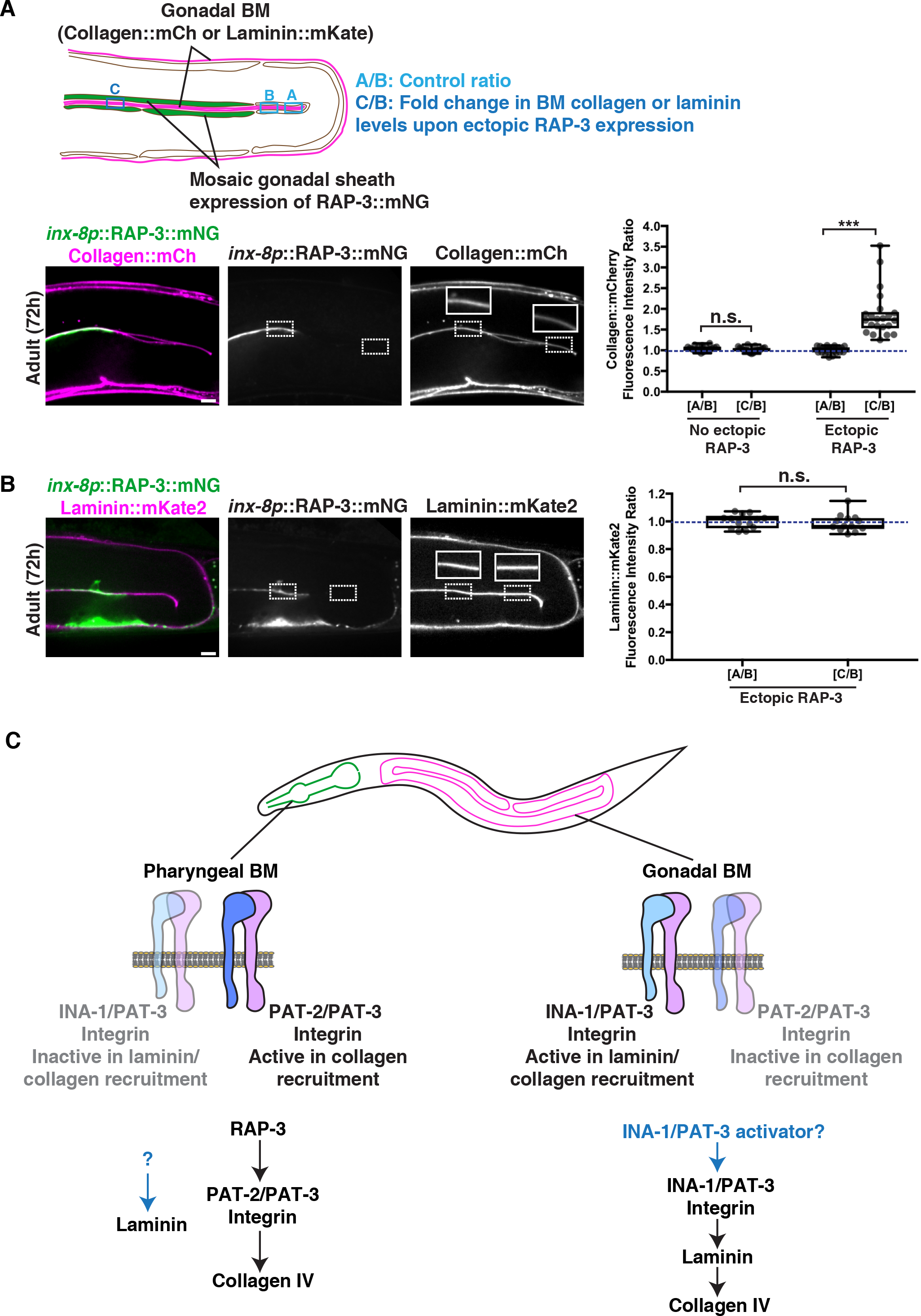
Ectopic gonadal expression of RAP-3 increases BM collagen but not laminin levels. **(A)** A schematic outlining mosaic expression of RAP-3∷mNG in the gonadal sheath cells and method for quantification of collagen and laminin levels is shown on top. A merged fluorescence image of *inx-8p*∷RAP-3∷mNG (green) and collagen∷mCh (magenta) in a 72h adult gonad is shown on the bottom left, and split RAP-3∷mNG and collagen∷mCh channel images on the bottom right. The dotted box on the left indicates a region of RAP-3 expression and the dotted box on the right denotes absence of RAP-3 expression. Collagen∷mCh signal in the dotted box regions are magnified in insets. Quantification of fold increase in gonadal BM collagen∷mCh fluorescence intensity upon ectopic RAP-3 expression is shown on the right (n=22). **(B)** A merged fluorescence image of *inx-8p*∷RAP-3∷mNG (green) and laminin∷mKate2 (magenta) in a 72h adult gonad is shown on the left, and split RAP-3∷mNG and laminin∷mKate2 channel images on the right. The dotted box on the left indicates a region of RAP-3 expression and the dotted box on the right denotes absence of RAP-3 expression. Laminin∷mKate2 signal in the dotted box regions are magnified in insets. Quantification of fold increase in gonadal BM laminin∷mKate2 fluorescence intensity upon ectopic RAP-3 expression is shown on the right (n=13). *** *p*<0.0001, paired two-tailed Student’s *t*-test; n.s., *p*>0.05. Box edges in boxplots depict the 25^th^ and 75^th^ percentile, the line in the box indicates the median value, and whiskers mark the minimum and maximum values. Scale bars are 10μm. **(C)** Model for distinct modes of collagen recruitment to the pharyngeal and gonadal BMs.

## Discussion

Type IV collagen is critical for BM function and numerous human diseases are associated with its misregulated accumulation or loss^7^. Previous work has suggested that a meshwork of laminin forms a cell-bound template that recruits the type IV collagen network through cross-bridging components^1,3^. How laminin is targeted to cell surfaces and whether there are other mechanisms to direct collagen to BMs *in vivo* has remained unclear. Through live-cell imaging of endogenous localization, conditional knockdown, misexpression, and RNAi screening, we have discovered distinct mechanisms for type IV collagen recruitment to the growing BMs of the *C. elegans* pharyngeal and gonadal tissues. We found that the two *C. elegans* integrin α subunits INA-1 (laminin-binding) and PAT-2 (RGD-binding) are expressed with the sole β subunit PAT-3 in both organs. Our results suggest each tissue promotes selective activation of specific integrin heterodimer to recruit collagen from the extracellular fluid: αINA-1/βPAT-3 activation in the gonad recruits laminin, which directs moderate levels of collagen to the BM; while αPAT-2/βPAT-3 activation in the pharynx recruits high levels of collagen independent of laminin. Supporting this model, we identified a putative pharyngeal-specific PAT-2/PAT-3 activator, the small GTPase RAP-3, an ortholog of mammalian Rap1 that mediates inside-out activation of mammalian integrins^49^. Collectively, these data reveal how tissues dictate collagen incorporation into BM through selective integrin activation and provide insight into how cells can use distinct mechanisms to target collagen to BMs, thereby precisely controlling collagen levels and constructing diverse BMs (Figure 8C).

Due to the challenge of imaging of BMs *in vivo*, the lethal phenotypes of null mutations of many BM components, and the expanded gene families of BM receptors and matrix components in vertebrates, it has been difficult to establish the mechanisms that mediate type IV collagen recruitment to BM *in vivo*^19,33^. Studies *in vitro* suggest several mechanisms might recruit laminin to cell surfaces, which in turn mediates type IV collagen incorporation. For example, work on embryoid bodies—groups of pluripotent embryonic stem cells—indicates that integrin β1 and dystroglycan matrix receptors function redundantly to promote the anchorage of laminin to the surfaces of cells^19^. Findings in cultured rat Schwann cells further suggests that sulfated glycolipids mediate laminin deposition^20^. Here, we exploited the small BM receptor and matrix families of *C. elegans*^6^, as well as their conditional knockdown by RNAi, to investigate how the sole *C. elegans* type IV collagen molecule is recruited to the BMs of growing gonadal and pharyngeal organs during larval development. We demonstrate that the two *C. elegans* integrin heterodimers play key roles in BM recruitment: αINA-1/βPAT-3 promotes laminin-dependent type IV collagen recruitment to the gonadal BM, while αPAT-2/βPAT-3 directs collagen IV to the pharyngeal BM independent of laminin. RNAi against these integrins did not reduce type IV collagen incorporation into the BMs as severely as knockdown of type IV collagen itself, suggesting that another cell surface system(s) may function with integrin. However, *C. elegans* does not synthesize sulfated glycolipids^35^, and neither knockdown of dystroglycan nor the matrix receptors glypican, discoidin domain receptors, LAR-RPTP, or teneurin reduced type IV collagen recruitment. Thus, integrins are likely the chief mediators of type IV collagen recruitment to the gonadal and pharyngeal BMs.

Laminin is required for the recruitment of type IV collagen to the BM of *Drosophila* and mouse embryos^9,17,18^. In addition, in embryoid bodies, the inhibition of laminin polymerization prevents the assembly of collagen IV on cell surfaces^50^. Further, in cultured Schwann cells, exogenously added collagen assembles into the BM-like matrix only in the presence of laminin^51^. Similarly, we found that collagen IV targeting to the gonadal BM was dependent on laminin. Comparatively little is known about laminin-independent collagen recruitment. BM collagen addition independent of laminin has recently been observed during wound repair in the *Drosophila* larval epidermal BM^52^, but the receptor(s) facilitating this mode of collagen targeting have not been identified. In our study, we found that the PAT-2/PAT-3 integrin is both required and sufficient for laminin-independent collagen IV recruitment to the pharyngeal BM. Sequence analysis suggests that PAT-2/PAT-3 is an RGD-binding integrin^53^. As multiple RGD sequences are present in both α1 and α2 chains of *C. elegans* collagen IV, it is possible that PAT-2/PAT-3 directly binds collagen, thus precluding the need for a laminin template. Supporting this notion, we found that mosaic over-expression of PAT-2/PAT-3 in the pharyngeal tissue caused an increase in BM collagen levels only in regions of the BM contacting overexpressed PAT-2/PAT-3. Countering this idea, it has been suggested that the RGD sequences in collagen may not be sterically accessible to integrins^54^. Hence, an alternative possibility is that PAT-2/PAT-3 binds either to a cell surface or BM molecule that directly binds to collagen.

Both the integrin α subunits INA-1 and PAT-2 and the β subunit PAT-3 are expressed in the gonad and the pharynx. Thus, tissue-specific integrin expression cannot account for the tissue-specific activity of gonadal INA-1/PAT-3 and pharyngeal PAT-2/PAT-3 in collagen recruitment. We considered the possibility that tissue-specific synthesis of laminin or type IV collagen might selectively activate gonadal INA-1/PAT-3 or pharyngeal PAT-2/PAT-3 heterodimers respectively as they are trafficked to cell surfaces (analogous to outside-in activation of integrin). However, this is unlikely as laminin is produced by both tissues^30^ and collagen is not synthesized in the pharyngeal epithelium^14^. Since the sole difference between the αINA-1/βPAT-3 and the αPAT-2/βPAT-3 integrins is the α subunit, we propose instead that tissue-specific factors activate αINA-1 or αPAT-2 to promote their tissue specific matrix-recruiting activity. One possible mode of activation is through the diverse cytoplasmic tail region of the α subunits. Studies in mammalian cell lines have shown that the cytoplasmic tails of α of integrin subunits are required for the inside-out activation of integrin receptors and may bestow specificity to integrin activation^38,55^. Supporting this idea, we found that expressing the PAT-2 integrin α cytoplasmic tail fused to the INA-1 extracellular domain in the pharynx activated its ability to recruit laminin, even though INA-1 does not normally function in pharyngeal BM laminin recruitment during larval development. One puzzling question is the functional significance of the α subunit expressed in each tissue and localized to the cell-BM interface, but not involved in BM recruitment. In the gonad, our evidence suggests that the α subunit PAT-2 acts in part to sequester the β subunit PAT-3, thus limiting the amount of active INA-1/PAT-3 heterodimer and thereby limiting laminin addition to the BM. It is also possible that α subunits not involved with BM recruitment might help localize proteases involved in matrix remodeling^56^ or might be associated with intracellular effectors that allow it to be activated in response to mechanical forces^57^.

In lymphocytes, co-expressed integrins αLβ2 and αMβ2, which share a common β subunit, are differentially activated through their divergent α cytoplasmic tails, triggering distinct adhesion responses to chemokine stimulation^58^. Notably, the mammalian small GTPase Rap1 facilitates the activation of the αLβ2 integrin through the lymphoid-enriched Rap1 effector molecule RapL, which binds to the αL subunit^59,60^. In our study, through a targeted RNAi screen of integrin-associated proteins, we identified RAP-3, a *C. elegans* ortholog of mammalian Rap1, as a pharyngeal-expressed activator of the PAT-2/PAT-3 integrin’s collagen IV recruiting function. *C. elegans* encodes two Rap1 orthologs, *rap-1* and *rap-3*^43^, with *rap-3* appearing to be more divergent^61^. We provide strong evidence that RAP-3 is a specific activator of PAT-2/PAT-3 integrin. First, similar to loss of PAT-2/PAT-3, loss of RAP-3 resulted in reduction of pharyngeal BM collagen but not laminin levels. Second, over-expression of a constitutively active mutant of RAP-3 in the pharyngeal epithelium increased collagen levels in regions of the pharyngeal BM contacting over-expressing cells, phenocopying our observations with pharyngeal PAT-2 over-expression. Third, our genetic analysis suggested that RAP-3 and PAT-2 act in the same pathway to promote collagen recruitment. Finally, ectopic expression of RAP-3 in the gonad increased type IV collagen levels in the gonadal BM, but not laminin levels, indicative of ectopic PAT-2/PAT3 activation. The effector(s) of RAP-3 that promotes PAT-2/PAT-3 activity in collagen recruitment to the BM is not known, but our data suggest it is likely expressed in both gonadal and pharyngeal tissues, as RAP-3 can activate PAT-2/PAT-3 in either tissue. Collectively, these observations support a model where the tissue-specific expression of RAP-3 in the pharynx promotes activation of PAT-2/PAT-3 to facilitate laminin-independent collagen recruitment to the BM, and suggests that distinct tissue-specific factor(s) might mediate INA-1/PAT-3 activation in the gonad to promote laminin-dependent BM collagen recruitment (Figure 8C).

The triple helical nature of collagen combined with its intermolecular covalent cross-links provides BMs their tensile strength, and genetic elimination of type IV collagen results in embryonic lethality in mice and *C. elegans* when tissues first experience mechanical loads^9,10^. Type IV collagen also tethers numerous matrix proteins and growth factors^7^. As a result of its complex and essential roles, precise levels of BM collagen are required for many cell and tissue functions. For example, the BMP/TGFβ ligand Dpp binds type IV collagen in *Drosophila* BMs, and collagen levels influence Dpp mediated signaling during dorsal ventral patterning in the embryo, germ stem cell production in the ovary, renal tubule morphogenesis, and wing disc growth^62–64^. Further, specific levels and gradients of type IV collagen mediate tissue constriction and shaping in the *Drosophila* egg chamber and wing disc and the *C. elegans* gonad^12,65,66^. Distinct integrin receptors that mediate different modes of type IV collagen recruitment may provide tissues robust mechanisms to control BM collagen levels. In particular, as PAT-2/PAT3 activity is controlled by the small GTPase RAP-3, its activity, and thus levels of type IV collagen, can be finely tuned through GTPase activating proteins (GAPs) and guanine nucleotide exchange factors (GEFs). This could be a dynamic mechanism for tissues to increase, decrease, and even translate signaling pathways into collagen gradients^12,67^. Therapeutically targeting specific activators of collagen-recruiting integrins may also be a promising means to modulate collagen levels during aging, in fibrotic diseases, and with type IV collagen genetic disorders where type IV collagen levels in BMs are altered, leading to tissue decline^7,68–70^.

## Acknowledgements

We thank Brent Hoffman, Laura Kelley, Kacy Gordon, David Reiner, Roy Zent, Thomas Hannich, Nick Brown, and members of the Sherwood laboratory for helpful discussions. Some strains were provided by the CGC, which is funded by NIH Office of Research Infrastructure Programs (P40 OD010440). E.L.H. was supported by postdoctoral fellowship 129351-PF-16-024-01-CSM from the American Cancer Society, and D.R.S. was supported by the National Institute of General Medical Sciences (Maximizing Investor’s Research Award R35 GM118049), and National Institute of Child Health and Human Development (R21 HD084290).

## Methods

### *C. elegans* strains

*C. elegans* strains used in this study are listed in Table S2. Worms were reared on NGM plates seeded with OP50 *Escherichia coli* at 16°C, 18°C, or 20°C according to standard procedures^71^.

### Transgene construction

#### mNeonGreen (mNG)/mKate2 knock-ins

To generate the following functional, genome-edited mNG knock-ins *lam-2(qy20[lam-2∷mNG + LoxP])*, *lam-2(qy41[lam-2∷mKate2 + LoxP])*, *pat-3(qy36[pat-3∷mNG + LoxP])*, *ina-1(qy23[ina-1∷mNG + LoxP])*, *pat-2(qy26[pat-2∷mNG + LoxP])*, *pat-2(qy49[pat-2∷2xmNG + LoxP])*, and *rap-3(qy57[rap-3∷mNG + LoxP])*, we used CRISPR/Cas9 genome editing with a self-excising drug (hygromycin) selection cassette (SEC) as described previously^72^, with some modifications. Briefly, we generated new starter repair plasmids lacking the 3xFlag tag downstream of the SEC cassette sequence, and we attached an 18 amino acid (glycine-alanine-serine) flexible linker (flexlink) in-frame and directly upstream of mNG, mKate2, or two tandem mNG fluorophores. ~700bp-1kb left and right homology arms for each target were amplified from N2 genomic DNA (gDNA), mutated to introduce six silent point mutations adjacent to the Cas9 cut site, and inserted into the appropriate repair plasmid using Gibson assembly. We generated 1 or 2 guide RNA (sgRNA) plasmids for each target by inserting the respective sgRNA sequences into the pDD122 plasmid. To direct cleavage near the C-terminus, the following sgRNA sequences were used: (5’-GTCATCAATTTGGAGCAAGA-3’) and (5’-ATTGATGACATTGAAGCATT-3’) for *lam-2*; (5’-TTGGCTTTTCCAGCGTATAC-3’) and (5’-TTTAAAAATCCAGTATACGC-3’) for *pat-3*; (5’-CGAGAAGAATGGGCTGATAC-3’) and (5’-ACGAGAAGAATGGGCTGATA-3’) for *ina-1*; (5’-CAGTACAATCAGGGACGTCA-3’) for *pat-2*; and (5’-ATCTAATCGTGTGCAAAACA-3’) for *rap-3*. For each tagging experiment, we injected a mixture of 50ng/μl Cas9-sgRNA plasmids (25ng/μl of each guide plasmid in cases where two sgRNAs were available), 100ng/μl repair template plasmid, and red co-injection markers (2.5ng/μl pCFJ90 (*myo-2p∷mCherry*) and 5ng/μl pCFJ104 (*myo-3p∷mCherry*)) into the gonads of ~30-40 young adult N2 animals. The injected animals were singled onto fresh OP50 plates and allowed to lay eggs for 3-4 days at 20°C in the absence of selection. Then, 500μl of 2mg/ml hygromycin solution was added to each plate, and the plates were returned to 20°C for 4-5 days. Candidate knock-in animals were roller [*sqt-1(e1350)*, dominant Rol mutation] worms that survived the hygromycin treatment and lacked the red fluorescent extrachromosomal array markers. We were able to isolate ~2-5 independent lines for each construct. All initial insertion strains were homozygous viable and segregated 100% Rol progeny. To excise the SEC, we heat-shocked plates containing ~6 L3/L4 rollers each at 34°C in a water bath for 4h, and then grew the animals at 20°C for 3-4 days. We then singled adult wild-type animals (worms that lost both copies of the SEC) and verified successful genome editing by visualizing mNG/mKate2 fluorescence, PCR genotyping, and sequencing of the fluorophore insertion site.

#### *rap-3* knock-out

To generate *rap-3(qy67[mNG + LoxP])*, we used CRISPR/Cas9 genome editing with the SEC as described above to delete the endogenous *rap-3* coding sequence and replace it with mNG. We directed cleavage near the N- and C-terminus of *rap-3* using the following sgRNAs: (5’-TTTTGGGAAATGGAGGAGTT-3’) and (5’-ATCTAATCGTGTGCAAAACA-3’). We modified the starter repair template plasmid described above to include 1kb left and right homology arms for *rap-3*.

#### myo-2p∷pat-2∷mNG∷unc-54 3’utr

*myo-2p∷pat-2∷mNG∷unc-54 3’utr* was built by Gibson assembling the following fragments in order: a ~1kb *myo-2p* fragment from the pCFJ90 vector, full-length *pat-2* amplified from N2 gDNA, and *flexlink∷mNG∷unc-54 3’utr*.

#### myo-2p∷ina-1[EX]∷pat-2[CTMD]∷mNG∷unc-54 3’utr

To generate *myo-2p∷ina-1[EX]∷pat-2[CTMD]∷mNG∷unc-54 3’utr*, we first amplified full-length *ina-1* and *pat-2* from N2 gDNA and cloned these fragments into separate TOPO vectors. Then we used Gibson assembly to ligate the following fragments in order: a ~1kb *myo-2p* fragment from the pCFJ90 vector; *ina-1* fragment containing only the extracellular, transmembrane, and intracellular membrane-proximal regions; *pat-2* fragment containing only the CTMD; and *flexlink∷mNG∷unc-54 3’utr*.

#### myo-2p∷rap-3^G12V^∷mNG∷unc-54 3’utr

To construct *myo-2p∷rap-3^G12V^∷mNG∷unc-54 3’utr*, we first used Gibson assembly to connect the following fragments in order: a ~1kb *myo-2p* fragment from the pCFJ90 vector, full-length *rap-3* amplified from N2 gDNA, and *flexlink∷mNG∷unc-54 3’utr*. We then introduced the G12V mutation in *rap-3* by site-directed mutagenesis.

#### inx-8p∷rap-3∷mNG∷unc-54 3’utr

To *build inx-8p∷rap-3∷mNG∷unc-54 3’utr*, we replaced (by Gibson assembly) the *myo-2p* fragment in the *myo-2p∷rap-3∷mNG∷unc-54 3’utr* construct created above with a 1.5kb fragment upstream of *inx-8* amplified from N2 gDNA.

### Transgenic strains

All CRISPR/Cas9 genome-edited mNG/mKate2 knock-in strains were created by injecting the relevant constructs into the germline of young adult N2 worms as described earlier. All knock-in alleles were functional, and viability and growth rates of knock-in strains were similar to N2 animals. The *rap-3(qy67)* knock-out strain was generated by injecting the relevant CRISPR/Cas9 constructs into the gonads of young adult NK364 animals. We were unable to isolate homozygous *rap-3(qy67)* animals as the null mutation caused embryonic lethality.

Transgenic worms expressing *myo-2p∷pat-2∷mNG∷unc-54 3’utr* or *myo-2p∷ina-1[EX]∷pat-2[CTMD]∷mNG∷unc-54 3’utr* were created by injecting these constructs (1.5ng/μl) together with full-length *pat-3* gDNA amplified from N2, EcoRI-digested salmon sperm DNA (50ng/μl), and pBluescript II (50ng/μl) into the gonads of young adult NK364 or NK2446 animals respectively. Animals expressing *myo-2p∷rap-3^G12V^∷mNG∷unc-54 3’utr* were created by co-injecting the construct (1.5ng/μl) with EcoRI-digested salmon sperm DNA (50ng/μl) and pBluescript II (50ng/μl) into the gonads of young adult NK364 animals. Worms carrying *inx-8p∷rap-3∷mNG∷unc-54 3’utr* were made by co-injecting the construct (50ng/μl), EcoRI-digested salmon sperm DNA (50ng/μl), pBluescript II (50ng/μl), and a green co-injection marker (*myo-2p∷gfp*, 2.5ng/μl) into the gonads of NK364 or NK2446 animals. As F1 progeny expressed the above constructs mosaically and did not transmit them to subsequent generations, we collected and imaged F1s directly for the relevant experiments.

### RNAi

All RNAi constructs were obtained from the Vidal^73^ and Ahringer^74^ libraries, except for the following clones: *csk-1*, *kin-32*, *pix-1*, *rsu-1*, *unc-97*, *unc-112*, and *pat-3*. To build these RNAi clones, we PCR-amplified fragments corresponding to the longest transcripts of these genes from N2 gDNA, and inserted them into the L4440 (pPD129.36) vector^75^ by Gibson assembly. RNAi constructs were then transformed into *Escherichia coli* strain HT115, and all RNAi experiments were performed using the feeding method^76^. Briefly, we grew RNAi bacterial cultures in selective media for 12-14h at 37°C, and then for an additional hour following addition of 1mM Isopropyl b-D-1-thiogalactopyranoside (IPTG) to induce dsRNA expression. For co-depletion experiments, we mixed the relevant induced bacterial cultures at a 1:1 ratio. NGM agar plates containing topically applied 1M IPTG and 100mg/ml ampicillin (9μl each) were then seeded with these RNAi bacterial cultures and left at room temperature overnight for further induction. Synchronized L1 worms were placed on RNAi plates and allowed to feed for 24-96h at 20°C, depending on the experiment. The L4440 empty vector was used as a negative control for all RNAi experiments. To improve RNAi knockdown efficiency in the pharyngeal tissue, we used worms harboring the *lin-35(n745)* mutant allele^77^; and for the gonadal sheath and other tissues, the *rrf-3(pk1426)* genetic background^78^. We verified knockdown efficiency for all RNAi experiments, with the exception of the targeted RNAi screen and *pat-2; pat-3* double RNAi experiments, by including a control with the relevant mNG/mCherry/mKate2-tagged target protein. We achieved between ~70-100% reduction for all RNAi experiments. For the co-depletion of PAT-2 and RAP-3, we verified knockdown efficiency by plate-level assessment of worm paralysis: RNAi initiated at the L1 larval stage against *pat-2* alone, or *pat-2* and *rap-3* in combination, resulted in 100% paralysis of animals by early adulthood.

### Imaging

Confocal DIC and fluorescence images were acquired at 20°C on an AxioImager A1 microscope (Carl Zeiss) controlled by μManager software^79^, and equipped with an EMCCD camera (Hamamatsu Photonics), a 40x Plan-Apochromat (1.4NA) objective, a spinning disc confocal scan head (CSU-10;Yokogawa Electric Corporation), and 488-nm and 561-nm laser lines. Worms were mounted on 5% noble agar pads containing 0.01M sodium azide for imaging. For all experiments except those detailed in Figure 1B and S3A, we captured single-slice images at the middle focal plane of animals where most or all of the gonadal and pharyngeal tissue cross-sections were in focus. For figure 1B, we acquired z-stacks at 1μm intervals spanning the entirety of the pharynx or gonad. For figure S3A, we captured single-slice images at a superficial focal plane, where the body wall muscle was in focus.

### Image analysis, processing, and quantification

All quantifications of mean fluorescence intensity were done on raw images in FIJI 2.0^80^. We drew ~5-8-pixel long linescans along the BM to obtain raw values of mean fluorescence intensity. Background intensity values were obtained by averaging two linescans of similar length in regions adjacent to the BM with no visible fluorescence signal. DIC and fluorescence images in all figures were processed in FIJI. The unsharp mask filter was applied to DIC images. 3D isosurface renderings of the pharyngeal and gonadal BMs were constructed using Imaris 7.4 software (Bitplane). Briefly, we acquired z-stacks of pharyngeal and gonadal collagen∷mCherry in early L1 and young adult worms. Young adult gonads were imaged in sections, and z-stacks of individual sections were stitched together with the FIJI pairwise stitching method^81^. 3D isosurfaces were constructed from these z-stacks by manually tracing the outline of BM collagen∷mCherry in every z-slice, and surface area measurements were automatically calculated from these isosurfaces by Imaris.

### Statistical analysis

Statistical analysis was performed in GraphPad Prism 7. To blind analysis of fluorescence intensity in all datasets, we used a filename-randomizing ImageJ macro (courtesy of Martin Höhne). Sample sizes were validated *a posteriori* through assessments of normality by log-transforming all datasets and then using the Shapiro-Wilk test. For comparisons of mean fluorescence intensities between two populations, we used an unpaired two-tailed Student’s *t*-test (with Welch’s correction in cases of unequal variance between samples). To compare mean fluorescence intensities between three or more populations, we performed one-way analysis of variance followed by either a post-hoc Dunnett’s or Tukey’s multiple comparison test. For comparisons of fluorescence intensity ratios, we used a paired two-tailed Student’s *t*-test (with Welch’s correction in cases of unequal variance between samples). Bar graphs and boxplots were prepared in GraphPad Prism 7. Figure legends indicate sample sizes, statistical tests used, and *p*-values.

---

**Figure S1.**
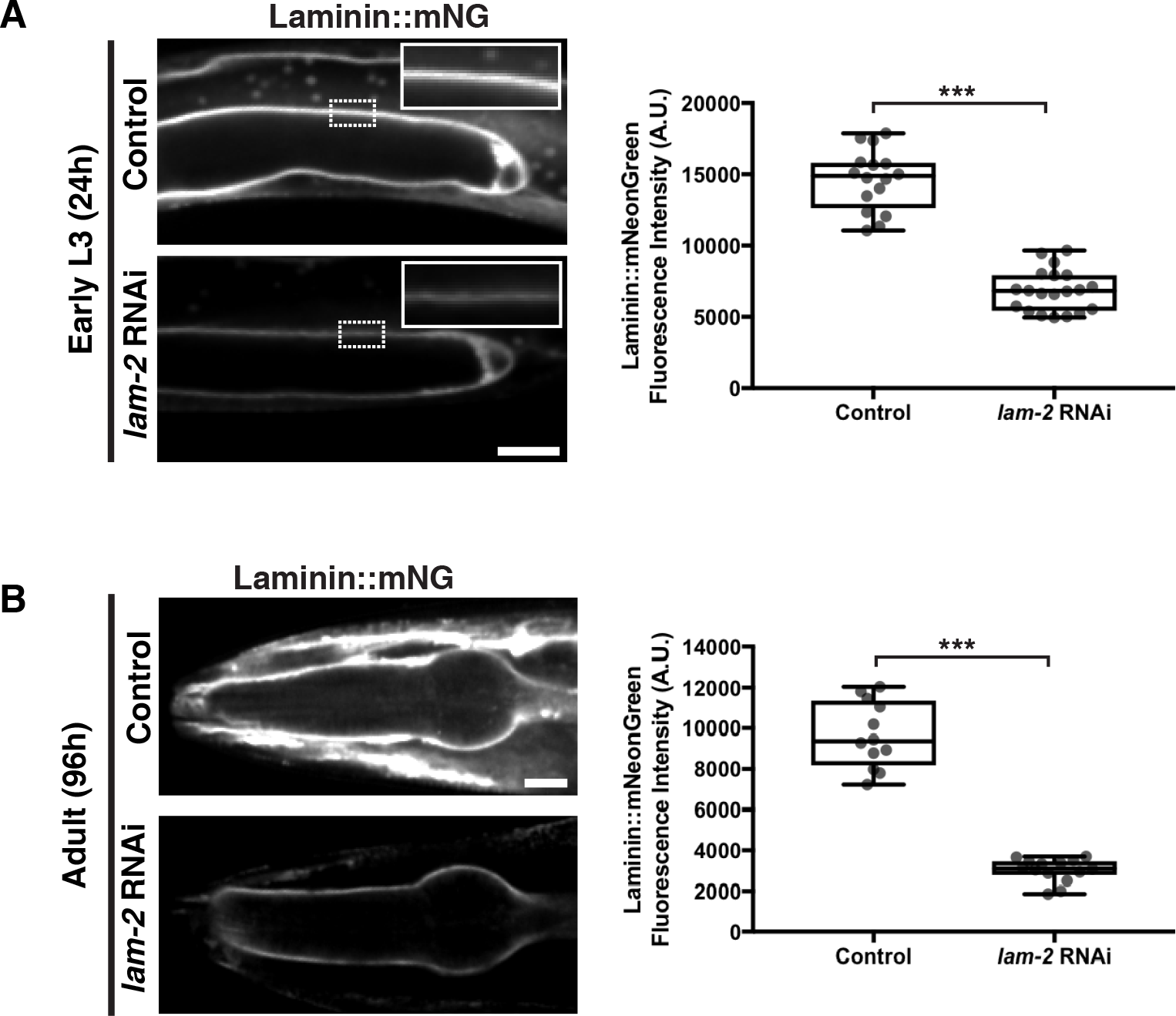
Pharyngeal and gonadal laminin levels after *lam-2* knockdown. **(A)** Fluorescence images of gonadal BM laminin∷mNG in control (L4440 empty vector) and *laminin* (*lam-2*) RNAi-treated animals (RNAi feeding initiated at the L1 stage and animals viewed at the early L3). Quantification of laminin∷mNG fluorescence intensity is shown in boxplots on the right (control n=16; *lam-2* RNAi n=20). **(B)** Fluorescence images of pharyngeal BM laminin∷mNG in control and *lam-2* RNAi-treated 96h adult animals. Quantification of laminin∷mNG fluorescence intensity is shown on the right (control n=12; *lam-2* RNAi n=14). \*\*\**p*<0.0001, unpaired two-tailed Student’s *t* test. Box edges in boxplots depict the 25^th^ and 75^th^ percentile, the line in the box indicates the median value, and whiskers mark the minimum and maximum values. Scale bars are 10μm.

**Figure S2.**
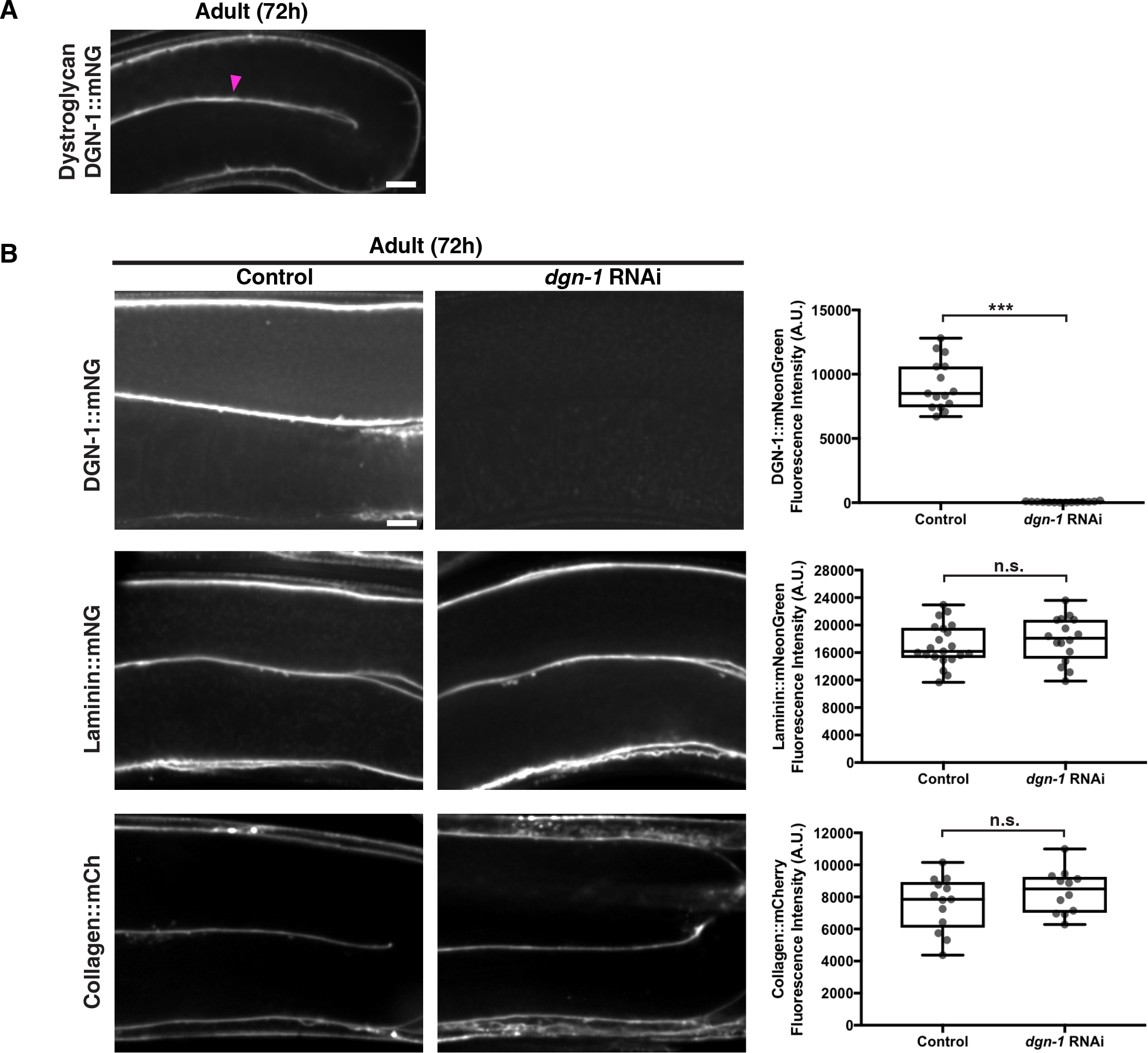
Dystroglycan is not required for laminin or collagen recruitment to the gonadal BM. **(A)** Fluorescence image of DGN-1∷mNG localization in a 72h adult gonad. The magenta arrowhead denotes enrichment of fluorescence signal at the cell-BM interface. **(B)** Fluorescence images of DGN-1∷mNG in the gonad in control and *dgn-1* RNAi-treated 72h adult animals is shown in the top panel, with quantification of DGN-1∷mNG fluorescence intensity on the right (control n=15; *dgn-1* RNAi n=14). The middle panel shows fluorescence images of gonadal BM laminin∷mNG in control and *dgn-1* RNAi-treated 72h adult animals, with quantification of laminin∷mNG fluorescence intensity on the right (control n=21; *dgn-1* RNAi n=16). The bottom panel shows fluorescence images of gonadal BM collagen∷mCh in control and *dgn-1* RNAi-treated 72h adult animals, with quantification of laminin∷mNG fluorescence intensity on the right (control n=13; *dgn-1* RNAi n=12). *** *p*<0.0001, unpaired two-tailed Student’s *t* test; n.s., *p*>0.05. Box edges in boxplots depict the 25^th^ and 75^th^ percentile, the line in the box indicates the median value, and whiskers mark the minimum and maximum values. Scale bars are 10μm.

**Figure S3.**
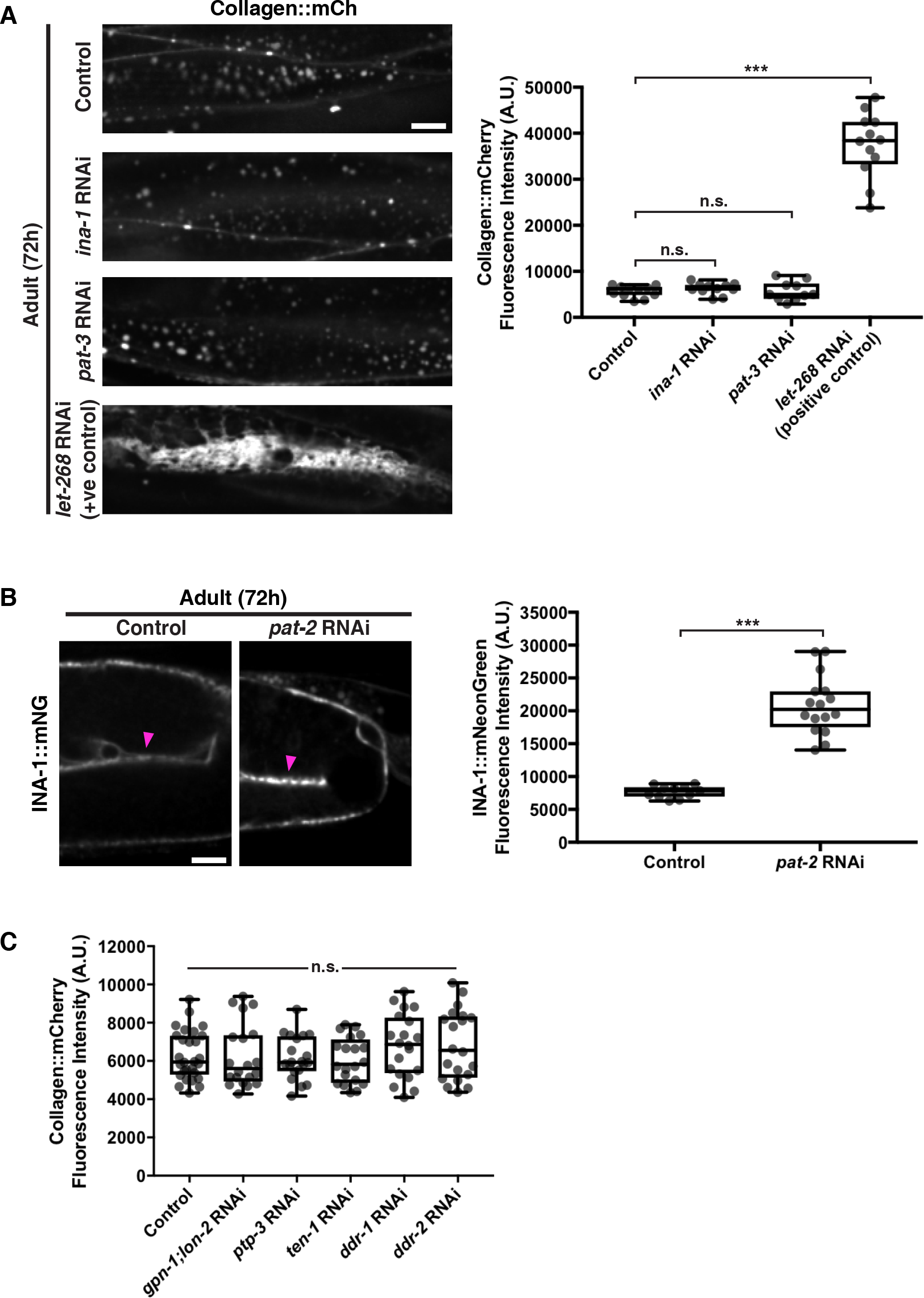
Knockdown of integrin does not perturb collagen secretion and integrin is the predominant receptor for gonadal BM collagen recruitment. **(A)** Fluorescence images of body wall muscle collagen∷mCh in control, *ina-1*, *pat-3*, and *let-268* RNAi-treated 72h adult animals, with quantification of collagen∷mCh fluorescence intensity on the right (control n=11; *ina-1* RNAi n=11; *pat-3* RNAi n=10; *let-268* RNAi n=12). *let-268* encodes a procollagen lysyl hydroxylase that is essential for collagen IV secretion^82^. *** *p*<0.0001, one-way ANOVA followed by post-hoc Dunnett’s test; n.s., *p*>0.05. **(B)** Fluorescence images of gonadal INA-1∷mNG in control and *pat-2* RNAi-treated 72h adult animals, with quantification of INA-1∷mNG fluorescence intensity on the right (control n=11; ina-1 RNAi n=16). The magenta arrowheads denote INA-1∷mNG enrichment at the gonad-BM interface. *** *p*<0.0001, unpaired two-tailed Student’s *t* test. **(C)** Quantification of gonadal BM collagen∷mCh fluorescence intensity in 72h adult animals upon knockdown of matrix receptors (control n=28 and n=20 each for all RNAi treatments). n.s., *p*>0.05, one-way ANOVA. Box edges in boxplots depict the 25^th^ and 75^th^ percentile, the line in the box indicates the median value, and whiskers mark the minimum and maximum values. Scale bars are 10μm.

**Figure S4.**
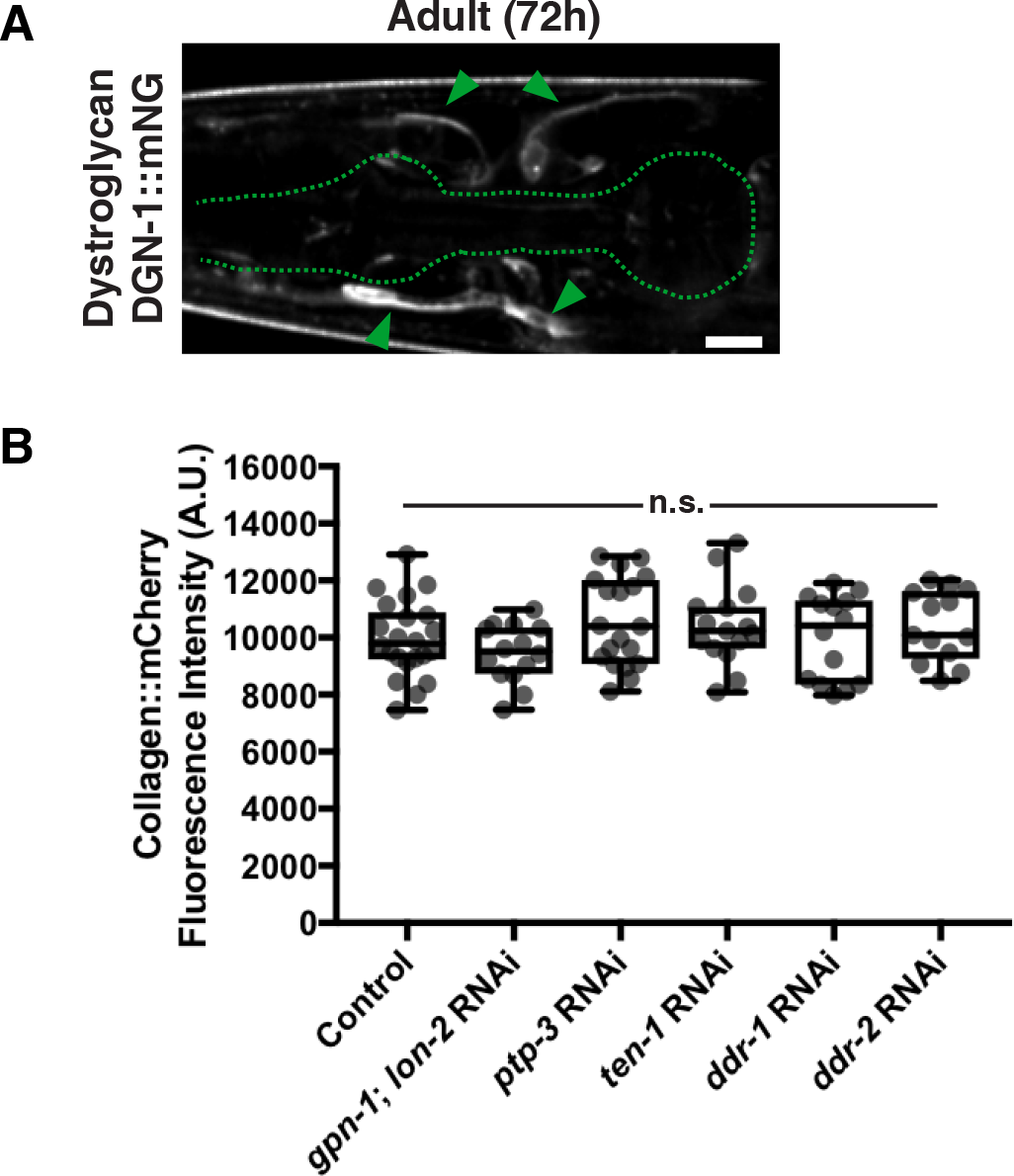
Dystroglycan is not expressed in the pharynx and PAT-2/PAT-3 is the predominant receptor for pharyngeal BM collagen recruitment. **(A)** Fluorescence image of DGN-1∷mNG localization in the head of a 72h adult animal. The pharynx is outlined in green. Green arrowheads indicate enrichment of DGN-1∷mNG at the nerve ring-nerve BM interface. Quantification of pharyngeal BM collagen∷mCh fluorescence intensity in 96h adult animals upon knockdown of matrix receptors (control n=22, *gpn-1, lon-2* RNAi n=14; *ptp-3* RNAi n=19; *ten-1* RNAi n=15; *ddr-1* RNAi n=14; *ddr-2* RNAi n=15). n.s., *p*>0.05, one-way ANOVA. Box edges in boxplots depict the 25^th^ and 75^th^ percentile, the line in the box indicates the median value, and whiskers mark the minimum and maximum values. Scale bars are 10μm.

**Figure S5.**
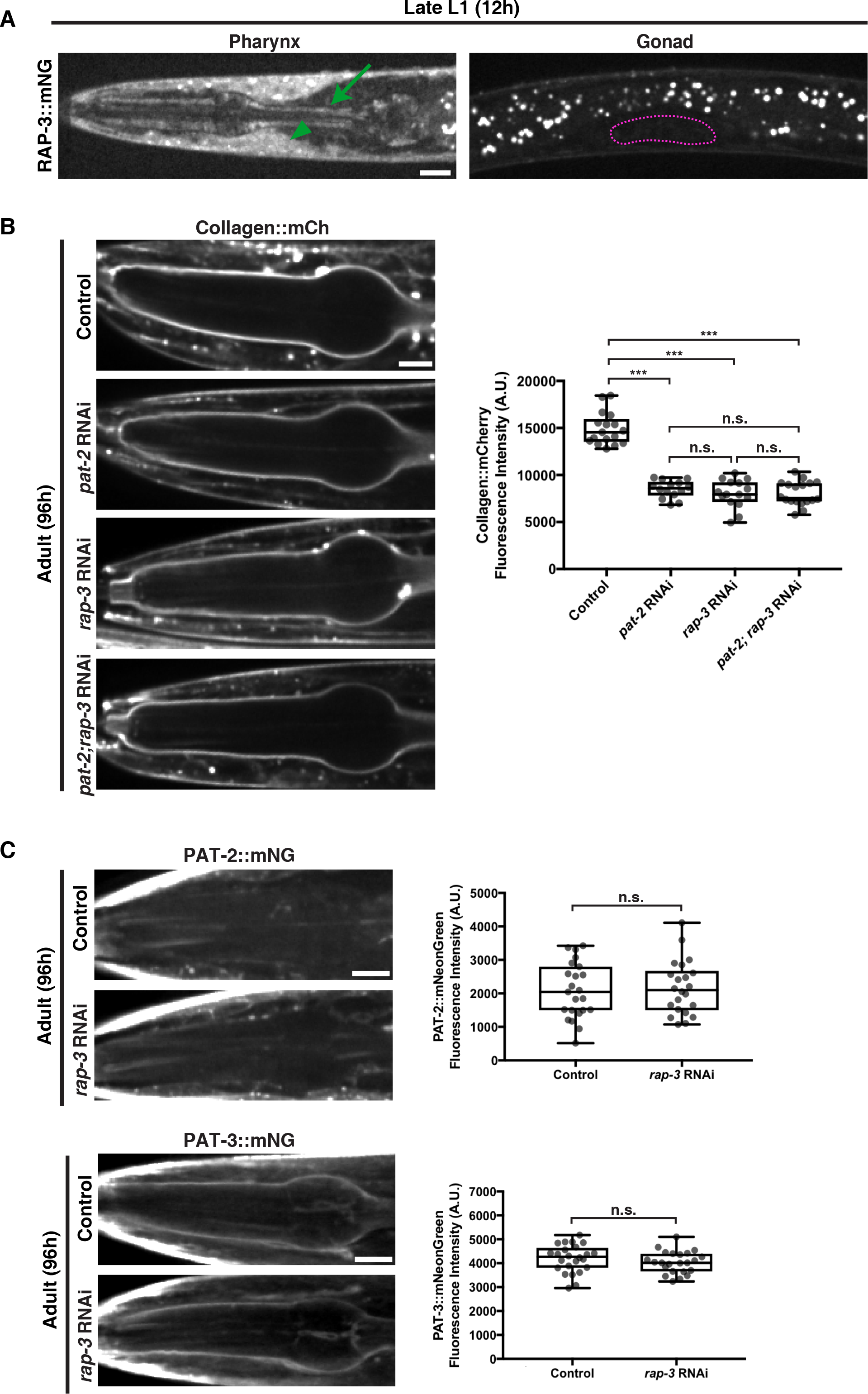
RAP-3 and PAT-2/PAT-3 integrin function together to promote pharyngeal BM collagen recruitment and RAP-3 does not regulate PAT-2 or PAT-3 levels. **(A)** Fluorescence images of RAP-3∷mNG localization in the pharynx (left) and gonad (right) of a late L1 animal. The green arrow indicates pharyngeal muscle localization of RAP-3∷mNG, while the green arrowhead denotes RAP-3∷mNG signal in the body wall muscle. The gonad is outlined in magenta. **(B)** Fluorescence images of pharyngeal BM collagen∷mCh in control, *pat-2*, *rap-3*, and *pat-2; rap-3* RNAi-treated 96h adult animals. Collagen∷mCh fluorescence intensity is quantified on the right (control n=17; *pat-2* RNAi n=14; *rap-3* RNAi n=15; *pat-2;rap-3* RNAi n=18). *** *p*<0.0001, one-way ANOVA followed by post-hoc Tukey’s test; n.s., *p*>0.05. **(C)** The top panel shows fluorescence images of pharyngeal PAT-2∷mNG in control and *rap-3* RNAi-treated 96h adult animals, with quantification of PAT-2∷mNG fluorescence intensity on the right (control n=23, *rap-3* RNAi n=22). The bottom panel shows fluorescence images of pharyngeal PAT-3∷mNG in control and *rap-3* RNAi-treated 96h adult animals, with quantification of PAT-3∷mNG fluorescence intensity on the right (control n=24, *rap-3* RNAi n=23). *** *p*<0.0001, unpaired two-tailed Student’s *t* test. Box edges in boxplots depict the 25^th^ and 75^th^ percentile, the line in the box indicates the median value, and whiskers mark the minimum and maximum values. Scale bars are 10μm.

**Table S1.**
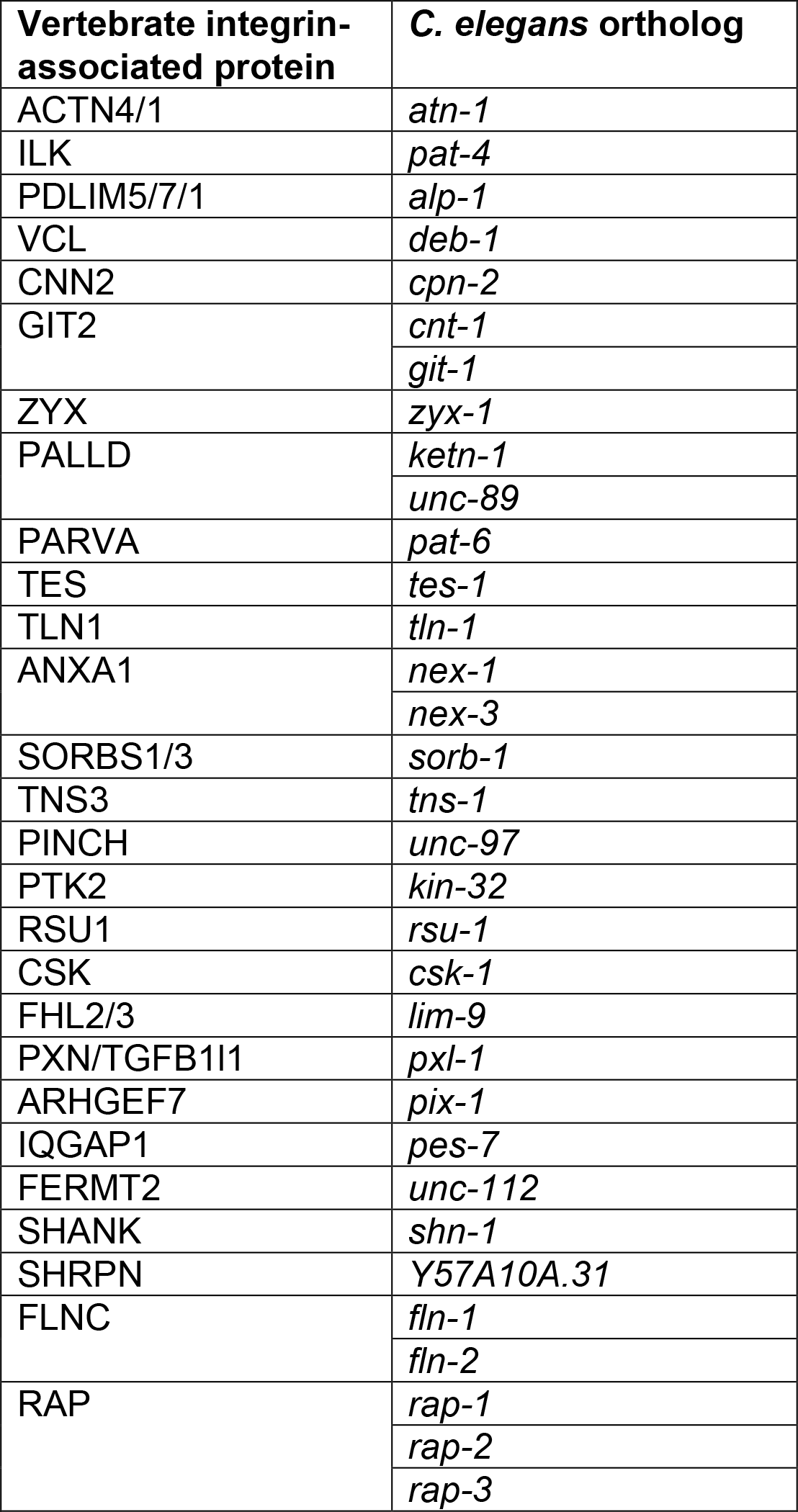
Targeted screen of vertebrate integrin-associated proteins with *C.elegans* orthologs.

**Table S2.** *C. elegans* strains.

| Strain | Genotype<sup>a</sup> | Reference |
| --- | --- | --- |
| N2 | Wild-type (ancestral) |  |
| NK364 | *unc-119(ed3)* III; **qyls46** [*emb-9p::emb-9::mCherry*, *unc-119(+)*] X | Ihara et al., 2011<sup>23</sup> |
| NK1911 | *rrf-3(pk1426)* II; *qyls46* X | This study |
| NK2335 | **qy20** [*lam-2::mNeonGreen*] X | This study |
| NK2351 | *lin-35(n745)* I; *rrf-3(pk1426)* II; **ruls38** [*myo-2p::gfp*, *unc-119 (+)*] III; *qyls46* X | This study |
| NK2334 | *rrf-3(pk1426)* II; *qy20* X | This study |
| NK2429 | *lin-35(n745)* I; *qy20* X | This study |
| NK2318 | **qy18** [*dgn-1::mNG*] X | This study |
| NK2469 | *rrf-3(pk1426)* II; *qy18* X | This study |
| NK2436 | **qy36** [*pat-3::mNG*] III | This study |
| NK2324 | **qy23** [*ina-1::mNG*] III | This study |
| NK2479 | **qy49** [*pat-2::2xmNG*] III | This study |
| NK2339 | *rrf-3(pk1426)* II; *qy23* III | This study |
| NK2064 | *rrf-3(pk1426)* II; *ruls38* III; *qyls46* X | This study |
| NK2446 | **qy41** [*lam-2::mKate2*] X | This study |
| NK2503 | **qy57** [*rap-3::mNG*] IV | This study |
| NK2538 | **qy67** [*rap-3 deletion + mNG*] IV; *qyls46* X | This study |
| NK2430 | *lin-35(n745)* I; **qy26** [*pat-2::mNG*] III | This study |
| NK2472 | *lin-35(n745)* I; *qy36* III | This study |

<sup>a</sup>When a transgene is listed for the first time, it is bolded and the entire genotype is displayed.

